# Neural signatures of motor memories emerge in neural network models

**DOI:** 10.1101/2025.04.02.646788

**Authors:** Joanna C. Chang, Claudia Clopath, Juan A. Gallego

## Abstract

Animals can learn and seamlessly perform a great number of behaviors. However, it is unclear how neural activity can accommodate new behaviors without interfering with those an animal has already acquired. Recent studies in monkeys performing motor and brain-computer interface (BCI) learning tasks have identified neural signatures—so-called “memory traces” and “uniform shifts”—that appear in the neural activity of a familiar task after learning a new task. Here we asked when these signatures arise and how they are related to continual learning. By modeling a BCI learning paradigm, we show that both signatures emerge naturally as a consequence of learning, without requiring a specific mechanism. In general, memory traces and uniform shifts reflected savings by capturing how information from different tasks coexisted in the same neural activity patterns. Yet, although the properties of these two different signatures were both indicative of savings, they were uncorrelated with each other. When we added contextual inputs that separated the activity for the different tasks, these signatures decreased even when savings were maintained, demonstrating the challenges of defining a clear relationship between neural activity changes and continual learning.

## Introduction

Animals can acquire, retain, and execute many different behaviors in a lifelong cycle in which new behaviors are learned without interfering with familiar ones. Learning leads to changes in synaptic connectivity ^1–4^ and neural activity ^5–12^. These changes can be retained on different timescales, from hours^13^ to decades^14^, and correlate with activity changes in different brain regions, from hippocampus^5^ to the basal ganglia^15^, cerebellum^7,8^, and the motor cortices^9–12,16^. While it is evident that learning new behaviors leads to new patterns in neural activity, it is unclear how brains can accommodate these new neural activity patterns with old ones. Specifically, how do brains integrate activity patterns that improve future performance of newly learned behaviors—a phenomenon called savings^17^ (Figure 1A)—without decreasing performance for older, familiar behaviors?

**Figure 1:**
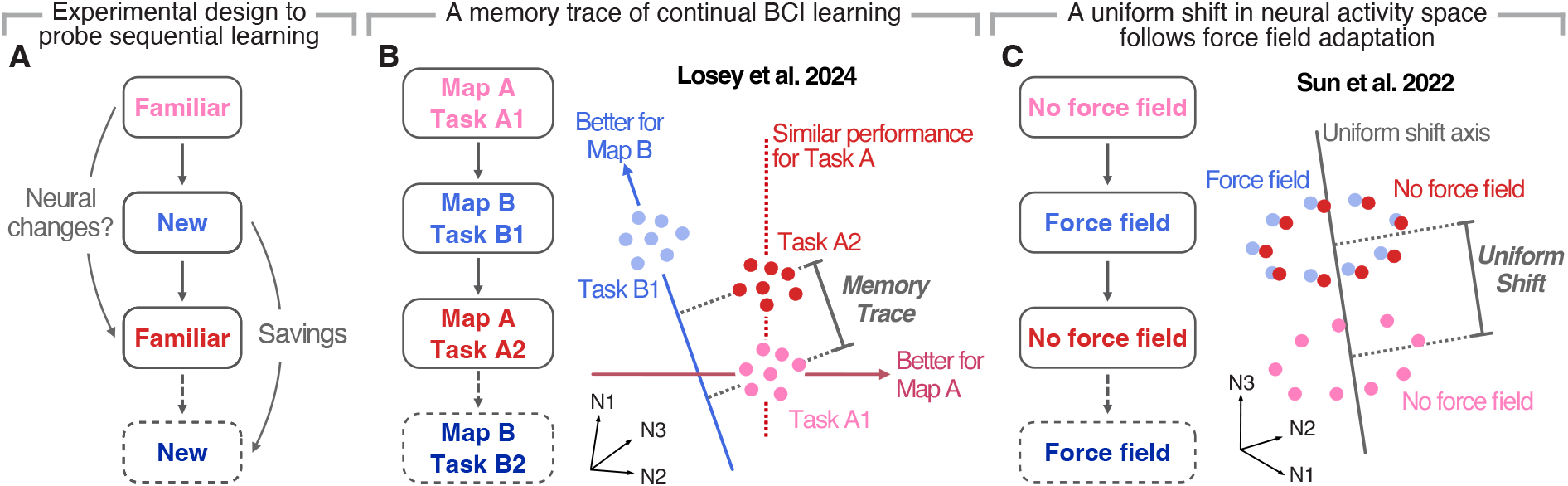
Potential neural signatures of learning. **A)** Learning a new task may cause changes in the neural activity used for a familiar task, and these signatures may help maintain knowledge about the new task to lead to retention savings if the new task is encountered again. **B)** Losey et al^18^ studied a BCI learning task where monkeys performed a center-out reaching task with a familiar Map A (Task A1, pink), learned a new Map B (Task B1, light blue), then returned to the familiar Map A (Task A2, red). They identified a memory trace that made neural activity (colored dots) during movement execution for the familiar Map A more useful for a new Map B after learning this new Map B. Importantly, the memory trace existed without compromising the performance for Map A. **C)** Sun et al^11^ studied a motor learning task where monkeys performed a baseline center-out reaching task (pink), learned to counteract a force field (light blue), then returned to the baseline task (red). They identified a uniform shift (colored dots) in the neural population space during movement preparation when monkeys learned to adapt to the force field that was sustained after returning to the baseline task.

Several recent studies have begun to elucidate how brains incorporate motor memories of new behaviors by adopting a neural population view ^11,18^. In this view, brain function is thought to be based on collective patterns of co-variation across a neural population rather than on the independent activity of its constituent neurons^19–21^. Since relatively few activity patterns are needed to explain the majority of variance in a neural population during a given task, neural population activity can be described mathematically based on a low-dimensional surface, a neural manifold^20^. Neural manifolds are thought to reflect biophysical (e.g., connectivity-related) constraints on neural activity ^10,22–24^ and have been leveraged to gain insights into the neural basis for learning^10,11,16,18,22^.

Adopting this view, Sadtler et al^22^ probed sequential learning using a brain-computer interface (BCI) paradigm where a linear map transformed neural population activity in monkey motor cortex to cursor movement, (Figure 1B). In each session, monkeys switched from a familiar Map A within their existing neural manifold (Task A1) to a new Map B (Task B1) and then back to the familiar Map A (Task A2). In different sessions, two different kinds of Map Bs were used to support the hypothesis that neural manifolds capture constraints on neural population activity: Map Bs that required activity patterns within an existing manifold (“within-manifold perturbations”) could be learned in a matter of minutes, whereas Map Bs that required activity outside the manifold (“outside-manifold perturbations”) could not be learned within a session, requiring progressive learning over several days^10^. Focusing on within-manifold perturbations in this BCI paradigm, Losey et al^18^ demonstrated that learning a new Map B altered the neural activity of the familiar Map A when monkeys were re-exposed to it (Task A2, Figure 1B). Crucially, this change made the activity during the familiar Map A more beneficial for the new Map B without negatively impacting performance of the familiar task (Figure 1B). The authors could show this directly by taking advantage of their BCI paradigm which, in contrast to traditional motor tasks where the relationship between motor cortex activity and behavior is not known, gave them full access to the relationship between neural activity and behavior since it is fully defined by the corresponding map: when they “projected” Task A2 activity onto Map B offline, it led to better performance for Map B than Task A1 activity. Thus, they proposed that Task A2 activity retained a “memory trace” of the new Map B that did not compromise the performance for the familiar Map A^18^.

Using a classic motor learning task in which participants—in this case, monkeys—performing reaching movements need to learn to counteract a force field^25^, Sun et al^11^ identified a “uniform shift” in the motor cortical population activity during movement preparation that emerged during learning. Similar to the “memory trace” in the previous BCI study, this shift was sustained after monkeys returned to the baseline task without a force field (Figure 1C). Since this shift was in an axis orthogonal to the directions in the neural manifold that were predictive of motor output, the authors argued that it could potentially index a motor memory of the new force field task, again in a way that would not compromise performance for the familiar baseline task^11^. Together, these BCI learning and motor learning studies suggest that learning new tasks may produce changes in neural activity that incorporate information about the new task in the activity of familiar tasks, potentially leading to savings if the new task is encountered again (Figure 1A). As a result, these signatures may be important for continual learning, that is, for learning tasks sequentially without forgetting previously acquired knowledge.

Here, we aimed to examine when these two potential signatures—the sustained uniform shift during preparation identified by Sun et al^11^ and the memory traces during execution identified by Losey et al^18^—arise and how they may relate to continual learning. To this end, we studied both signatures during the same learning paradigm using recurrent neural networks (RNNs). We used RNNs because similar models have been shown to replicate key features of neural activity and motor output from experimental recordings^26–30^, enabling investigation of scenarios that are difficult to probe experimentally^26,31,32^.

We modeled an adapted version of the sequential learning BCI task described above, which allowed us to compare both the behavior and the changes in network activity—including the two potential signatures of motor memories we are interested in—to those observed in monkey motor cortex in the original studies. We found that both sustained uniform shifts and memory traces emerged naturally in our RNNs, without modeling any explicit additional mechanisms to force them to arise. To relate these signatures to continual learning, we examined a common way to measure the extent of continual learning: savings, or improved relearning of a task upon re-exposure, compared to the initial exposure ^17^. Savings can be observed as either an increase in performance due to better retention immediately after re-exposure (“retention savings”), or a faster rate of learning during prolonged exposure (“learning rate savings”). Here, we focused on retention savings since learning rate savings might be confounded by different initial errors for our models trained with gradient descent. While the magnitude of the memory traces were closely correlated with retention savings, that of the uniform shift was not correlated with neither the magnitude of the memory traces nor the retention savings. Extending Losey et al’s ^18^ analyses to outside-manifold perturbations, we saw that memory traces were larger for outside-manifold perturbations when compared to within-manifold perturbations. This was due to greater forgetting in within-manifold perturbations when reverting back to the familiar map in Task A2. Inspired by behavioral studies on context-dependent learning ^33,34^, we added a context cue to better separate activity patterns and reduce interference to prevent forgetting. With context cues, a uniform shift was observed during learning of the new map in Task B1, but it did not persist during Task A2. Similarly, the magnitude of memory traces decreased and was no longer indicative of savings, even though there was still increased savings of the new map. Therefore, uniform shifts and memory traces arise naturally when learning a new task that shares activity patterns with a familiar task, but these signatures of learning for a novel task are no longer observed in the activity of the familiar task if they are separated by features like strong context cues.

## Results

### An RNN model of sequential learning naturally recapitulates experimental results

To understand how memory traces^18^ and uniform shifts^11^ arise and to explore their features in the context of continual learning, we used RNNs to model the same BCI experiments as in Losey et al^18^ (Figure 2A). In these experiments, monkeys used their motor cortical activity to control a computer cursor through a BCI map that translated neural activity onto cursor velocity. To study how neural activity changed across different tasks, monkeys had to control the BCI to produce center-out reaches using different BCI maps that were presented in a sequential block design.

**Figure 2:**
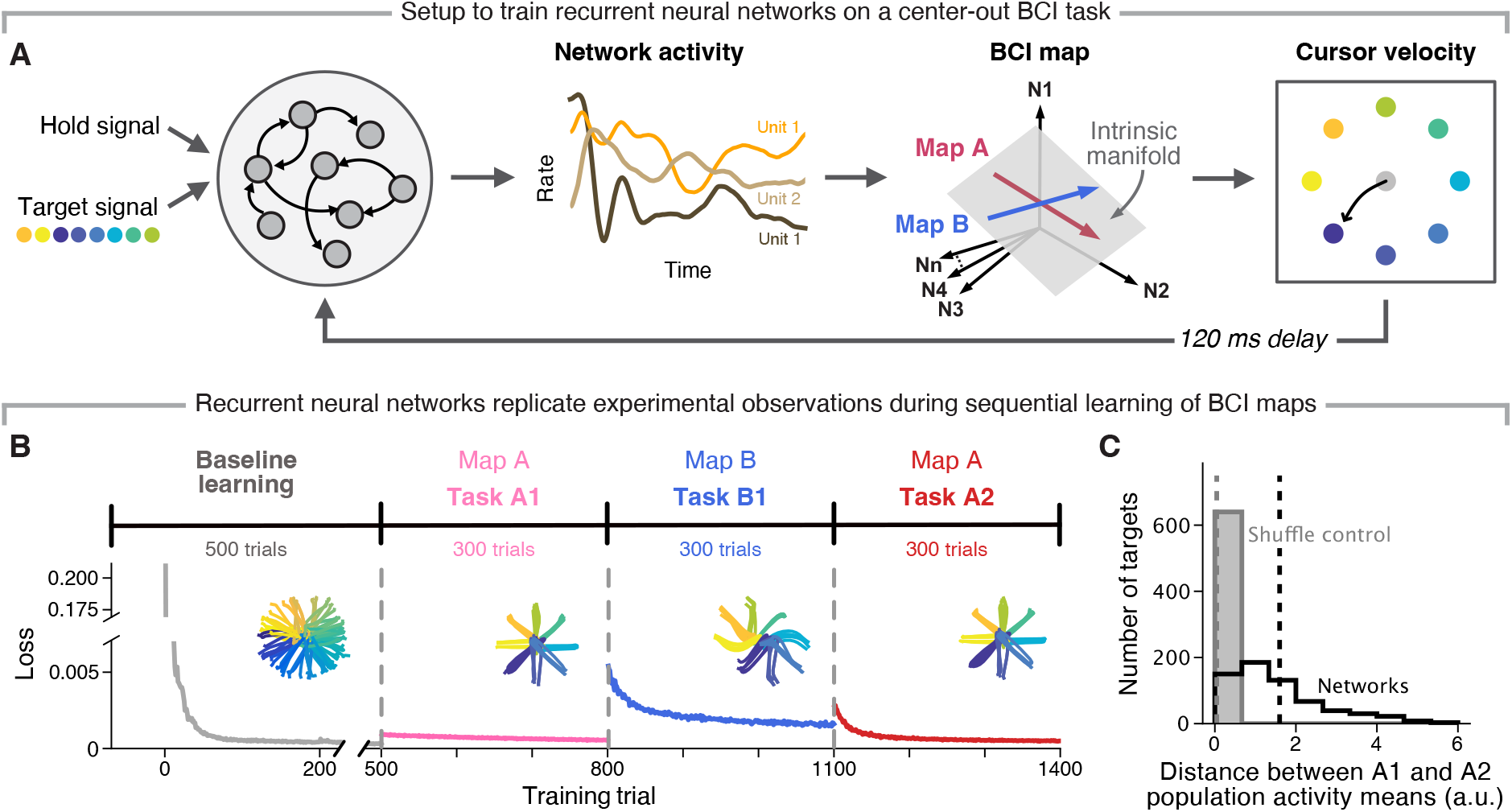
RNNs recapitulate experimental results in sequential learning. **A)** Networks were trained to perform center-out reaches by using different BCI maps which linearly transformed network activity to output velocities. Map A was the intuitive map based on the “intrinsic manifold” –a low-dimensional flat surface that captured the dominant covariation patterns of the network activity after baseline learning–, while Map B was a permutation of Map A. Produced velocities were fed back into the network after a 120 ms delay. **B)** Sequential training procedure: we first trained the recurrent, input, and output network weights during an initial training phase where networks had to produce random center-out reaches. Then, networks were trained to produce center-out reaches to eight fixed targets using different BCI maps (A or B) as a fixed output layer. Networks learned to produce output through these maps in blocks: from Map A (Task A1), to Map B (Task B1), then back to Map A (Task A2). Loss curves during each task block for an example network. Inset: position trajectories at the end of each block. **C)** Mahalanobis distance between population activity means for Task A1 and Task A2 for each target (black), and for controls where the task labels were shuffled (gray). Dotted lines, means across 10 different maps for each random seed (*n*=8 random seeds).

Here, we first used gradient descent to pretrain RNNs to produce velocities for varied delayed center-out reaches to model the monkeys’ ability to produce the necessary neural activity for controlling their arms (Figure 2B). We incorporated a delay period to examine possible uniform shifts during preparation, even if the original BCI experiments did not have one. RNNs were also given delayed feedback of their produced velocities^26^ (Figure 2A), and their performance was quantified as the mean-squared error (MSE) between the produced and synthetic target velocities (Methods). Following this initial training, RNNs were trained to perform the eight-target center-out BCI task using different BCI maps, which were modeled as fixed output weights. Each BCI map consisted of two transformations. First, activity from all 400 units in our RNNs was projected onto an eight-dimensional flat neural manifold, which we identified using Principal Component Analysis (PCA) and thus captured the majority (89-91%) of the variance in the network activity. Second, activity within this manifold was projected to the two-dimensional subspace defined by the corresponding BCI map to obtain the output cursor velocity along the horizontal and vertical axes (Figure 2A).

To probe whether and how learning a new task affected the neural activity of a familiar task, the RNNs were trained on the two different maps in sequential blocks: first with Map A (Task A1), then with a new Map B (Task B1), then back to the familiar Map A (Task A2, Figure 2B). Replicating the experiments^22^, Map A was determined based on the “intuitive manifold” map that captured the majority of the variance in the activity following baseline training, and Map B was a within-manifold perturbation of Map A that required activity in the same low-dimensional neural manifold (Methods).

The RNNs were able to learn Map A and Map B in sequence, with similar rates of learning and performance as the experimental results^18,22^ (Figure 2B). Recapitulating experimental results, neural activity patterns for a given target changed from Task A1 to Task A2 in our models, even if the mapping during Tasks A1 and A2 was the same^18^ (Figure 2C, Figure S1A). These changes were significant across different targets and significantly greater than a shuffled control (Figure 2C, *P* = 8.9 *·* 10^−107^, one-sided Wilcoxon signed-rank test). Thus, learning a new task (even a new version of the same task, in this case defined by Map B) altered the RNN activity for a familiar task without the need for an additional learning mechanism.

### Uniform shifts and memory traces arise from sequential learning without adding explicit mechanisms

Examining the properties of these activity changes revealed that they could be partially attributed to a uniform shift in the preparatory population activity from Task A1 to Task B1, which was orthogonal to the directions in neural space predictive of motor output (Figure 3A, Methods), as experimentally observed in monkey motor cortex^11^. These activity shifts generally persisted in Task A2 (Figure 3A-C; Figure S2A,B), further replicating experimental results^11^. Due to these changes, the different blocked tasks (Task A1, Task B1, and Task A2), could be easily distinguished based on their preparatory unit activity alone using linear discriminant analysis classifiers (Figure S1A), demonstrating that the tasks were indeed using different activity patterns.

**Figure 3:**
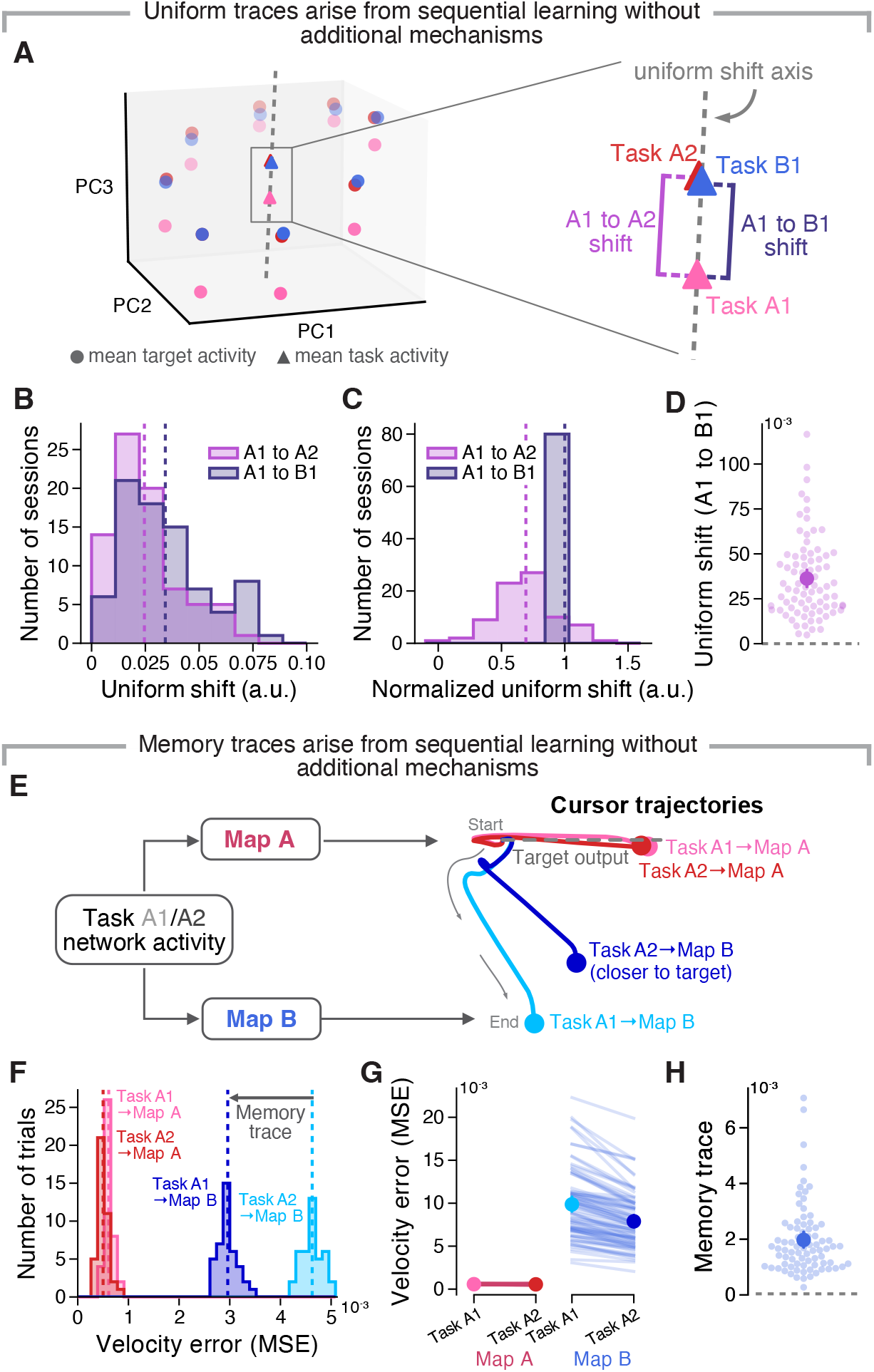
Uniform shifts and memory traces arise from sequential learning without explicit mechanisms. **A)** Population activity during preparation (first 200 ms of each trial) for an example session. Circles, time-averaged, trial-averaged activity for each target (mean target activity) in each task block. Triangles, time-, trial-, and target-averaged activity (i.e., mean task activity) in each block. The uniform shift axis is quantified by the direction of the vector connecting the mean task activity from Task A1 to Task B1. The uniform shift across tasks is calculated by projecting the mean task activity onto the uniform shift axis and calculating the resulting distance between tasks. **B)** Uniform shift from Task A1 to Task B1 and Task A1 to Task A2. Dotted lines, means across 10 different maps for each random seed. **C)** Normalized uniform shift, calculated by dividing by the uniform shift from Task A1 to Task B1 for each session. **D)** Uniform shift from Task A1 to Task B1 across targets. Big circle and error bars, means and 95% confidence intervals with bootstrapping for 10 different maps for each random seed (*n*=8 random seeds); small circles, different maps. Note that uniform shifts are consistently greater than zero (dotted line) indicating a shift in population activity. **E)** Task A1 and Task A2 activity were projected into Map A online or into Map B offline. The resulting output was used to assess how useful the activity is for each map, i.e., how close to the target output it brings the cursor. **F)** MSE between the produced and target output for each projection for an example target in an example session. Dashed lines, mean MSE across trials for that target. The memory trace is quantified as the decrease in mean MSE (i.e., the increase in performance) from Task A1**→**Map B to Task A2**→**Map B. **G)** MSE across all targets for projections onto the two different maps. Circles and error bars, means and 95% confidence intervals with bootstrapping for 10 different maps for each random seed (*n*=8 random seeds). Traces, different sessions. **H)** Same as D but for the memory trace. Note that memory traces are consistently greater than zero (dotted line) indicating increased offline performance for Task B with activity used for Task A2 compared to that for Task A1.

Having established that uniform shifts emerge naturally as a result of continual learning in RNNs using standard training techniques, we next asked whether we could observe similar memory traces as those found in experimental recordings^35^. Specifically, we were interested in whether memory traces spontaneously arise from our simple sequential learning model, or if an additional explicit mechanism was necessary for them to arise.

The advantage of a BCI setup is that we can probe how useful the neural activity for a familiar map is for a new BCI map before and after learning the new BCI map. While Task A1 and Task A2 used the same Map A online, we could project their activity through Map B offline to evaluate how much the activity changed from Task A1 to Task A2 to accommodate new activity patterns that were directly useful for Map B. In other words, we asked how much learning Map B left a memory trace of this new task in the activity of the familiar task defined by Map A (Figure 3E). When projected through Map A, Task A1 and Task A2 activity both produced output position trajectories that were similar to the target output, without a clear difference in performance (example in Figure 3E). However, when projected through Map B, Task A2 activity produced output trajectories that were closer to the target output than that of Task A1 activity (Figure 3E), indicating the existence of a memory trace of Map B. We quantified this memory trace as the increase in performance, or decrease in MSE, for Map B from Task A1 to Task A2 (Figure 3F). From here, we will denote “Task X activity projected through Map Y” as “Task X**→**Map Y”, such that the memory trace is the decrease in MSE from Task A1**→**Map B to Task A2**→**Map B.

We observed that memory traces consistently arose across different targets (Figure 3G,H; Figure S3A). These increases in performance for Map B from Task A1 to Task A2 were significantly greater than a control based on the performance for Map A (Figure 3G, *P* = 3.9 *·* 10^−15^, one-sided Wilcoxon signed-rank test), demonstrating that the presence of these memory traces did not result in a trade-off or loss of performance for Map A. This indicates that information from both maps coexisted in the activity of the familiar task as RNNs performed Task A2. Importantly, these memory traces could be specifically attributed to the process of learning Map B: when we reset the input and recurrent network weights after Task B or randomly reallocated the weight changes that occurred during Task B, the memory traces disappeared (Figure S3B,C). As a result, our models showed that both sustained uniform shifts and memory traces can arise spontaneously through sequential learning, potentially allowing the networks to fit both tasks in their activity repertoire even without any added mechanisms to reduce interference.

### The magnitude of memory traces, but not of uniform shifts, correlates with savings and initial behavioral error

Having established that our RNN models readily reproduce potential neural signatures of motor memories observed during BCI learning^18^ and motor learning^11^ experiments, we asked whether these signatures have a clear relationship with behavior.

First, we studied whether they captured the formation of a motor memory. Behaviorally, exposure to a perturbation leads to the formation of a motor memory, which is expressed by increased task performance upon re-exposure. One way motor memories manifest is through retention savings, defined as the increase in task performance immediately after (re-)exposure. We quantified these retention savings by comparing performance when networks were re-exposed to Map B during a new Task B2 compared to initial exposure to this same map during Task B1. Interestingly, the magnitude of the memory trace was closely related (Figure S4A) and highly correlated with the magnitude of the savings (Figure 4A, Figure S5D), but the magnitude of the uniform shift was not (Figure 4C, Figure 4I). Thus, even if both signatures spontaneously arose during continual learning, their relationship with behavior is different, and only the memory trace seems to be related to immediate retention savings.

**Figure 4:**
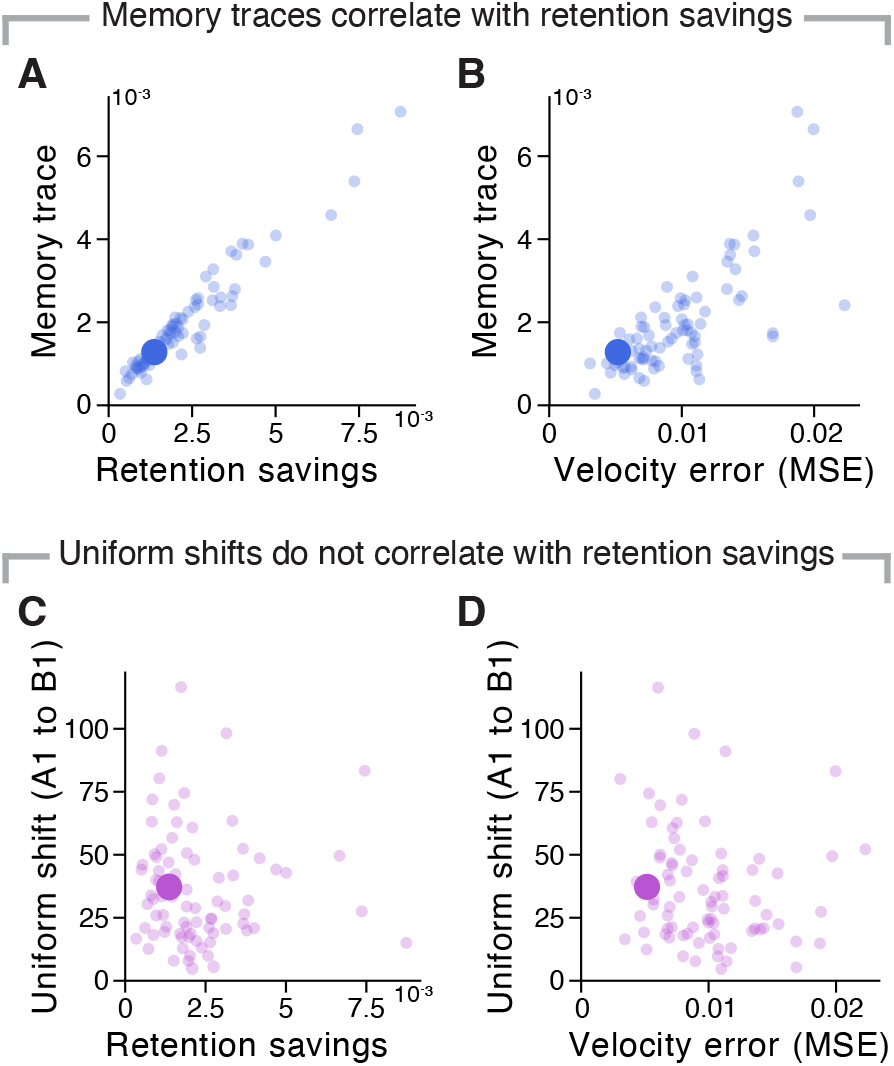
The magnitude of memory traces but not that of uniform shifts correlates with retention savings. **A)** Mean memory trace across targets compared to the retention savings. Small circles, mean for each map (*n*=10 maps) for each random seed (*n*=8 random seeds); large circle, example shared across A-D. **B)** Mean memory trace across targets compared to the initial velocity error (MSE) when Task A1 activity is projected into the new Map Bs before learning in Task B1 (i.e. Task A1**→**Map B MSE). **C)** Uniform shift from Task A1 to Task B1 across targets compared to the retention savings. **D)** Uniform shift from Task A1 to Task B1 across targets compared to the initial velocity error (MSE).

Having defined these relationships, we asked whether we could predict their magnitude from task properties alone. The BCI learning paradigm, combined with our modeling setup that allows us to run many “experiments”, offers a unique opportunity to answer this question. We characterized the Map B perturbations based on the directions of the neural manifold they affected (Figure S5A), since recent theoretical work suggests that large variance directions may be easier to control than their lower variance counterparts^36^. While “high variance perturbations” led to larger memory traces than “low variance perturbations” (Figure S5B), such difference in variance did not affect the magnitude of the uniform shifts (Figure S5C). Other measures such as the principal angles between BCI maps, or their relative amount of neural variance explained also had inexistent or weak associations with uniform shifts and memory traces (Figure S5E,F,G,J,K,L). In contrast, the initial behavioral disruption imposed by the perturbation was a strong predictor of the magnitude of the memory trace (Figure 4B, Figure S5H), although it did not predict that of the uniform shift (Figure 4D, Figure S5M). This suggests that more information is retained about a given task as measured by the memory trace if the task requires more learning at the behavioral level.

### Experimentally hard-to-learn outside-manifold perturbations lead to larger memory traces and smaller uniform shifts

We have studied uniform shifts and memory traces for BCI maps that impose within-manifold perturbations that require new activity patterns within the existing neural manifold. Here, we extended our investigation to outside-manifold perturbations that require activity patterns that are not available within the existing manifold (Figure 5A). Behaviorally, these perturbations are very different: while within-manifold perturbations can be learned within a few minutes ^22^, outside-manifold perturbations cannot, requiring several days ^10^.

**Figure 5:**
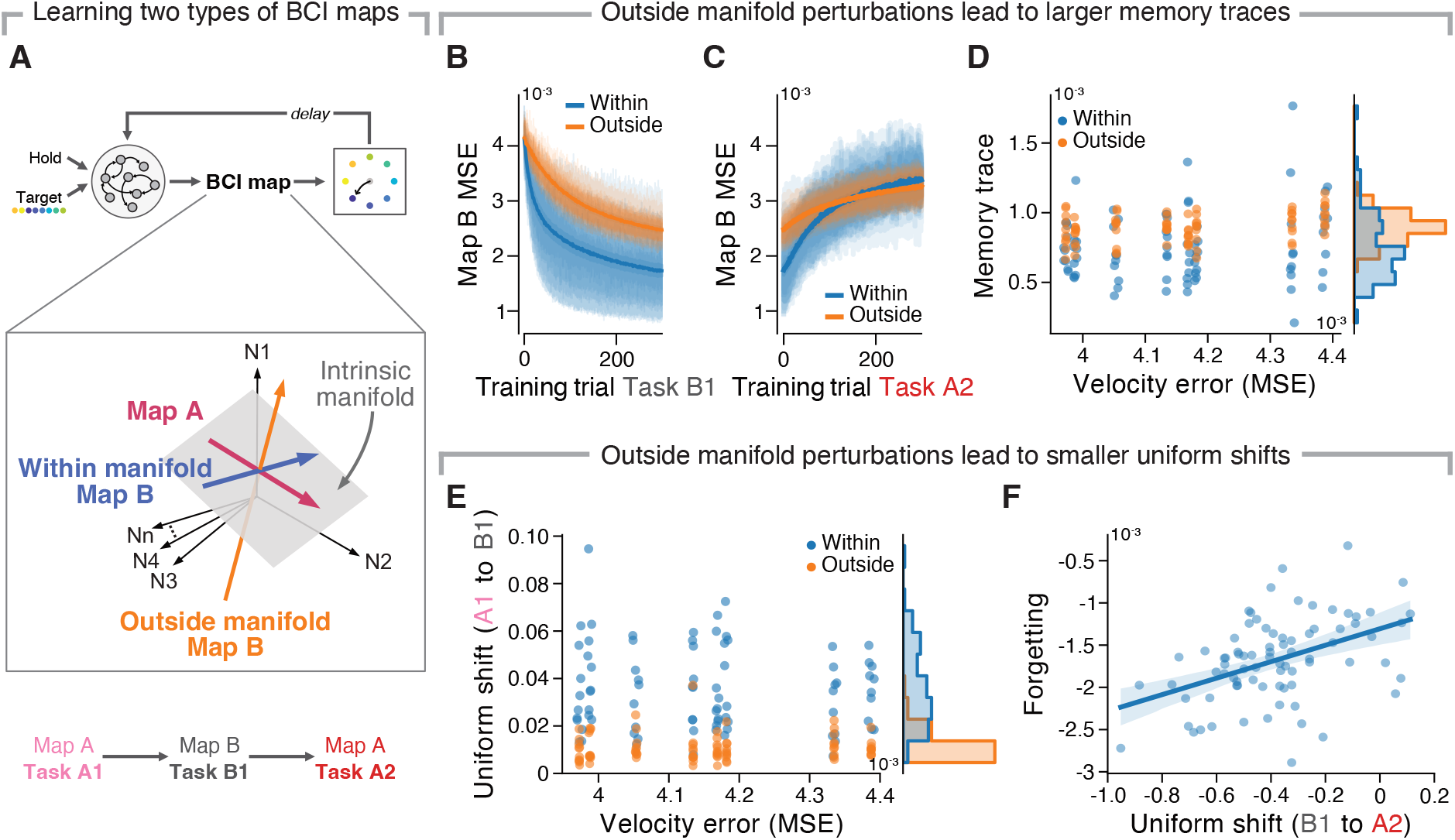
Learning new outside-manifold maps leads to larger memory traces than learning new within-manifold maps due to less forgetting. **A)** Networks were trained on different Map Bs representing different within- and outside-manifold perturbations that had the same Task A1**→**Map B MSE. **B)** Map B MSE during initial learning of the new Map B in Task B1. Lines and shaded surfaces, mean and 95% confidence interval across 10 maps for each perturbation across networks of different seeds (*n*=8 random seeds). Lines, different maps for different seeds. **C)** Map B MSE during reversion to the familiar Map A in Task A2. **D)** Mean memory trace across targets compared to the Task A1**→**Map B MSE. Circles, mean for each map for each random seed. Note that points with the same Task A1**→**Map B MSE belong to the same seed since this was controlled for. **E)** Uniform shift from Task A1 to Task B1 across targets compared to the Task A1**→**Map B MSE. **F)** Uniform shift from Task B1 to Task A2 compared to forgetting, defined as the decrease in performance from Task B1**→**Map B to Task A2**→**Map B, for within-manifold perturbations. Lines and shaded surfaces, mean and 95% confidence interval across different maps for different seeds; circles, different maps for different seeds.

Having established that the initial task error was a strong predictor of memory trace magnitude for within-manifold perturbations, we asked how uniform shifts and memory traces change when we control for this factor for both inside- and outside-manifold perturbations (Figure 5A). To do so, we projected Task A1 activity into thousands of within-manifold perturbation and outside-manifold perturbation BCI maps and selected a subset of maps of each type with the same MSE to use as the new Map Bs (*n* = 10 for each perturbation type, Methods, Figure S6A). Recapitulating experimental findings ^22^, outside-manifold perturbations were harder to learn than within-manifold perturbations, with larger errors at the end of Task B1 when the same number of training trials were used to learn the maps (Figure 5B, *P* = 8.2*·*10^−30^, one-sided Student’s t-test).

Learning outside-manifold maps less well would suggest smaller initial retention savings of these maps compared to within-manifold maps. However, when we examined the memory traces at the end of Task A2, outside-manifold perturbation led to surprisingly larger memory traces and savings than within-manifold maps (Figure 5D, *P* = 2.3 *·* 10^−7^, one-sided Mann-Whitney U test; Figure S6C, Figure S7A).

To reconcile these findings, we examined performance for the new maps during re-exposure to the original map (i.e., Task A2) and saw that the performance for within-manifold maps dropped more quickly than that for outside-manifold maps (Figure 5C). This suggests that during Task A2, information about within-manifold maps was more easily overwritten and forgotten, likely because these maps were too similar to coexist together without interference. To directly examine this, we trained networks initialized from the same random seeds to learn both Map A and Map B simultaneously right after initialization. RNNs had more difficulty learning two within-manifold maps simultaneously than one within-manifold and one outside-manifold map, showing that it was harder for within-manifold maps to coexist (Figure S7B). This supports our interpretation that the larger memory trace after learning an outside-manifold perturbation mapping happened because, although these maps were learned less well in the first place, more information about the new maps was retained after switching back to the familiar task (Figure 5C), leading to greater retention savings. The greater savings during outside-manifold perturbations were again correlated with greater memory traces (Figure S6B; note that this effect did not result from a tradeoff in performance for the familiar map, Figure S8). In contrast, our second potential neural signature of motor memory exhibited the opposite behavior: the magnitude of the uniform shift from Task A1 to Task B1 was smaller for outside-manifold perturbations than within-manifold perturbations (Figure 5E, *P* = 1.6 *·* 10^−22^, one-sided Mann-Whitney U test; Figure S2E,F), and was not directly correlated with retention savings (Figure S6H). However, the degree of forgetting of Map B was correlated with retraction along the uniform shift axis during Task A2 for within-manifold perturbations (Figure 5F), showing that, the degree of retention of the uniform shift, rather than its absolute length, was indicative of savings. Together, these results show that more similar tasks, like within-manifold perturbation maps, may be initially easier to learn, but may also cause greater interference and less retention during continual learning.

### Explicit context cues alter memory traces, uniform shifts, and their relationship to savings

Since forgetting due to interference was a driving factor in reducing savings for within-manifold perturbations, we examined ways to reduce interference to promote continual learning. Thus far, we have not introduced any explicit mechanisms to separate the activity patterns for different tasks. However, experimental work shows that humans and monkeys are better at learning tasks in blocks than when they are interleaved^11,37,38^, a phenomenon that is likely due to the incorporation of contextual information^33,34,38^. Based on this, we predicted that explicitly cuing our RNNs on the identity of the task would allow different task-specific activity patterns to be learned sequentially without overwriting. This intuition was built on the notion that an explicit cue signal would promote activity patterns for the different tasks to emerge in different parts of activity space.

To test this intuition, we added explicit context cues as inputs to our networks that indicated whether they were controlling the familiar Map A or the new Map B using one-hot encoded binary vectors (Figure 6A). We trained networks on the same sequential learning setup as before, but now with this additional context cue: Context A was used during baseline learning, Task A1, and Task A2, while Context B was used during Task B1. Using this setup, we asked how continual learning was modulated by the degree of discriminability between contexts—since such discriminability will likely indicate the degree that activity is driven to occupy different parts of state space—, and how this would translate into differences in uniform shifts and memory traces. We controlled the degree of discriminability between contexts by multiplying the binary vectors representing the context (Map A or Map B) by different factors to create context cues of varying magnitudes, and separately trained networks with each context magnitude.

**Figure 6:**
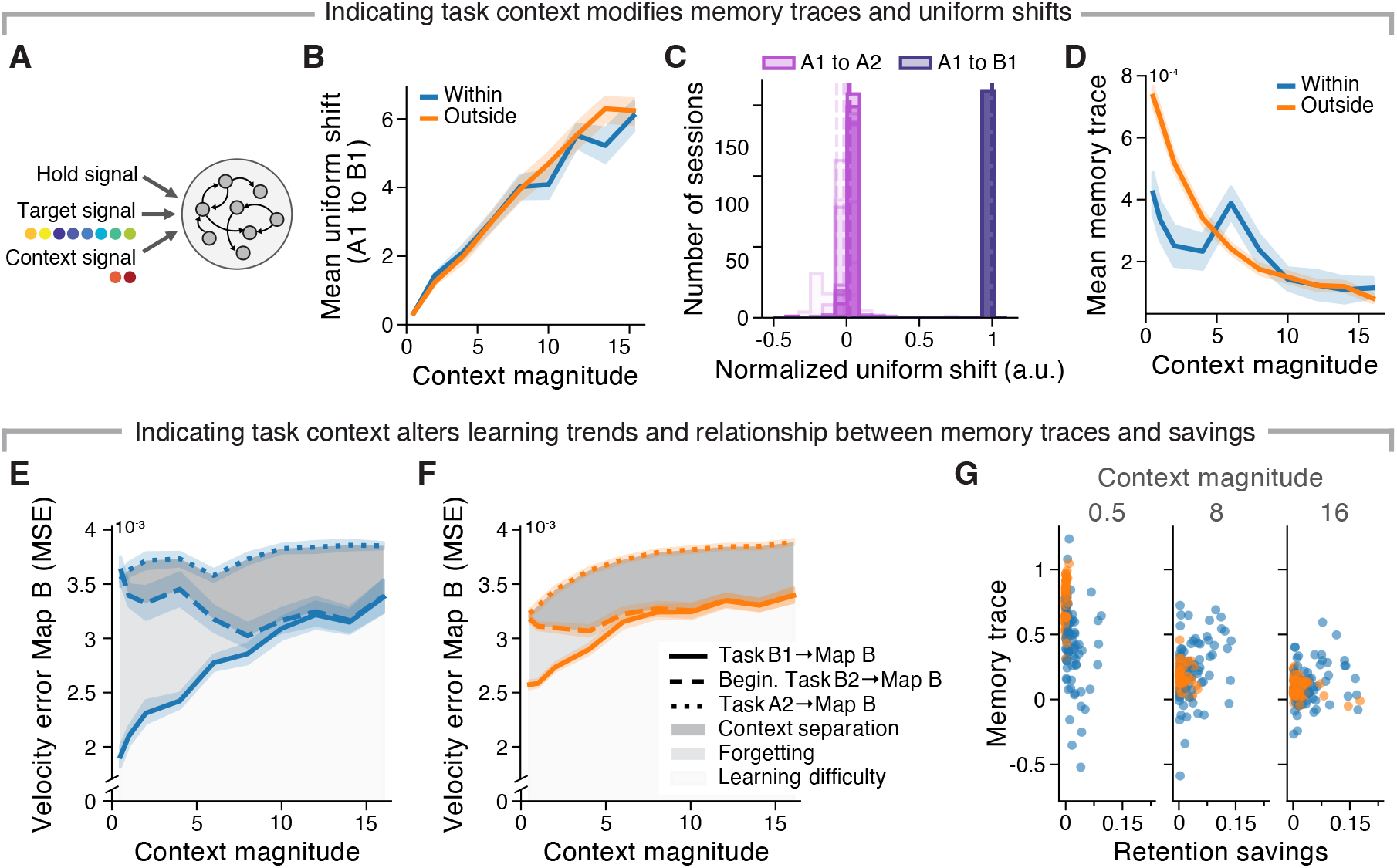
Context cues alter uniform shifts and memory traces. **A)** Networks were given an additional context cue that signaled whether the familiar Map A or the new Map B was being used. The context signal was a one-hot encoded vector, and the vector was multiplied by different factors to create different context magnitudes. **B)** Uniform shifts from Task A1 to Task B1 for different context magnitudes. Line and shaded surfaces, mean and 95% confidence interval across 10 maps for each perturbation across networks of different seeds (*n*=8 random seeds). **C)** Normalized uniform shifts from Task A1 to Task B1 and from Task A1 to Task A2, averaged across targets for each session. Shifts were normalized by the length of the shift from Task A1 to Task B1. Opacities, different context magnitudes (most transparent for smallest magnitude). Dashed lines, mean for each context magnitude. **D)** Memory trace for different context magnitudes. Line and shaded surfaces, mean and 95% confidence interval across maps for each perturbation across networks of different seeds. **E)** MSE when activity from the end of Task B1, the beginning of Task B2, and the end of Task A2 are projected into Map B for different context magnitudes for within-manifold perturbations. Task B1**→**Map B MSE indicates learning difficulty, the difference between Task B1**→**Map B and beginning of Task B2**→**Map B MSE indicates forgetting, and the difference between the beginning of Task B2**→**Map B and Task B1**→**Map B MSE indicates activity changes due to context. **F)** Same as E but for outside-manifold perturbations. **G)** Mean memory trace across targets compared to the retention savings. Small circles, mean for each map for each random seed.

As a result of introducing context, the trends in uniform shifts and memory traces that we previously saw dramatically changed. First, uniform shifts were still evident when learning the new map, and the magnitude of the shifts increased with greater context magnitudes (Figure 6B, Figure S2G,Figure S9A). However, the uniform shifts were not sustained: upon returning to the familiar Map A in Task A2, the uniform shift disappeared (Figure 6C, Figure S2G). This suggests that context signals pushed activity for the new map to occupy different spaces from the familiar map, but these changes were not incorporated into activity for the familiar map. Thus, context can lead to a dramatic change in this signature of learning.

Second, the inclusion of context cues also changed the trends in memory traces. Initially, with small context magnitudes, the trends were similar to those in networks without any context cues (Figure 5): outside-manifold maps led to greater memory traces than within-manifold maps (Figure 6D). Moreover, even though within-manifold perturbations were easier to learn during Task B1, there was also greater forgetting during Task A2, leading to greater Task A2**→**Map B MSE and smaller memory traces overall^1^ (Figure 6E,F).

As the context magnitude increased, however, the magnitude of the memory traces became indistinguishable for outside-manifold perturbation and within-manifold maps (Figure 6D). This trend was caused by larger context magnitudes leading to more disparate activity patterns which occupied a different part of state space ((Figure S10), making learning new maps more difficult. With less learning (Figure 6E,F), there was less knowledge about the new map that could be retained, contributing to a general decrease in memory traces with larger context magnitudes (Figure 6 E,F). With activity patterns in a different part of activity space due to the increased context magnitude, there was also less overwriting and forgetting of Map B during Task A2 (Figure 6E,F). Thus, greater context magnitudes made within-manifold perturbations more similar to outside-manifold perturbations: both were equally difficult to learn and equally resistant to forgetting, with essentially perfect retention savings (Figure 6E,F). However, unlike our previous networks without context cues, here the savings were no longer closely correlated with the memory trace (Figure 6G, Figure S9, Figure S4A). With context however, the context cue shifted Task B2 into a different part of state space from Task A2, such that Task A2 activity became less relevant for Map B (Figure S4A). This context separation made larger context magnitudes lead to smaller memory traces of Task B even when there was perfect retention savings. Together, these results show that there can still be motor memories of a new task that lead to savings even when the uniform shift is no longer sustained or if the memory trace is decreased. With little or no context, memory traces were good indicators for initial retention savings and the degree of retention of the uniform shift was a good indicator for forgetting. However, these relationships deteriorated when a large context input was added.

## Discussion

Using RNNs, we showed that learning a new task naturally causes activity changes that are observed when returning to a familiar task, recapitulating results from several experimental learning studies in monkeys ^11,18,22^. Learning new tasks both caused uniform shifts in the preparatory activity ^11^ that were sustained when reverting to a familiar task, and left memory traces that made activity for the familiar task more appropriate for the new task during movement execution^18^. These changes appeared naturally, without the addition of an additional mechanism, and allowed information about both tasks to coexist in the same neural activity patterns, facilitating retention savings. However, when we added contextual inputs that separated the activity for the different tasks, both the retention of the uniform shift and the memory trace were reduced, showing that these signatures of motor memory may not be present even if savings are maintained.

### Relationship to continual learning

Our results show that uniform shifts and memory traces indeed are signatures for continual learning, but have certain caveats. To measure the extent of continual learning, we have focused on the retention savings, the increase in performance immediately after re-exposure due to greater retention. While Sun et al^11^ specifically correlated the magnitude of the uniform shift during relearning to learning rate savings, Losey et al^18^ did not directly relate memory traces to any measures of savings. Here, we focused on retention savings since learning rate savings might be confounded by starting at different motor output errors at the beginning of re-exposure (Task B2) for our models trained with gradient descent. In our models, the magnitude of memory traces was closely correlated with retention savings following both within-manifold and outside-manifold perturbations (Figure 4A, Figure S4A, Figure S5D) for networks without contextual inputs, while the magnitude of uniform shifts was not (Figure 4D, Figure S5I). Rather, uniform shifts were indirectly related to retention savings: the relative retention of the uniform shifts was correlated with the amount of forgetting (Figure 5F), which affected the amount of savings for within-manifold perturbations (Figure 5C). Thus, our simulations and analyses show that while memory traces and uniform shifts could both be related to continual learning, they are uncorrelated with each other (Figure S6B).

Importantly, while these measures could be useful signatures for continual learning, they are not always indicative of savings, and they are not necessary for continual learning to occur. By including context signals, we pushed activity to occupy different parts of neural activity space, leading to a decrease in memory traces and a lack of retention of uniform shifts (Figure 6C,D). This suggests that if context inputs or other “top-down” inputs drive tasks to use very different activity patterns for producing motor output across different tasks—in the extreme case, pushing different tasks to be performed by non-overlapping subpopulations of neurons—, these signatures can disappear even if motor memories of new tasks are retained. Here, our RNN specifically modeled neural activity in motor cortex which is directly used for BCI control, so such inputs may originate from activity upstream of motor cortex. However, motor learning and adaptation involve contributions from many regions other than motor cortex, including parietal cortex^39,40^, cerebellum^41–43^, and basal ganglia^44,45^. While learning was constrained to our RNN model of motor cortex for the majority of this work, we also examined results when we added a second upstream RNN module and constrained learning to happen in this module. Even when learning was restricted to this upstream module, memory traces and sustained uniform shifts still arose (Figure S11), suggesting that the presence of these phenomena does not necessitate learning in the area that produces the behavioral output, in this case the motor cortex.

### Effects of context on within-manifold and outside-manifold perturbation learning

By modeling the BCI paradigm from Sadtler et al^22^ with many different within- and outside-manifold perturbations, we were able to further examine the differences between these perturbations. Previous work has largely distinguished the two classes of perturbations based on factors such as learning timescales^22^, neural changes required for learning^10,46^, reliance on correct feedback^27^, and modeled synaptic weight changes^47^. Here, we showed that there was variability in the memory traces and uniform shifts within each class of perturbations, highlighting differences between different perturbations in each class. While outside-manifold perturbations generally led to larger memory traces, some still overlapped with the distribution of within-manifold perturbations, suggesting that the changes that within- and outside-manifold perturbations impose on neural activity may live on a continuum, rather than fall within two discrete categories. Moreover, adding and increasing the magnitude of a context signal led to less forgetting for within-manifold perturbations (Figure 6E), causing them to adopt similar results as outside-manifold perturbations (Figure 6D,E,F).

These unexpected similarities between within- and outside-manifold perturbations in the presence of large context cues were likely caused by these cues pushing supposedly within-manifold perturbations to be outside the estimated manifold, as the original manifold estimated during baseline learning did not capture activity patterns imposed by different contexts. This is supported by large decreases in the activity’s overlap with the manifold when the context switched at the beginning of Task B, even before learning occurred (Figure S10). Previous work has shown that neural manifolds are generally nonlinear^48^, so large upstream inputs like context signals may drive activity into a part of activity space that is not captured by the same flat manifold estimated using a linear dimensionality reduction method—as we have done here and in the original experimental work. Thus, depending on the magnitude of these signals, we may explore activity that is more or less within the estimated flat manifold, rather than distinctly fully within or outside the actual nonlinear manifold. Indeed, this continuum between within- and outside-manifold perturbations may explain why even in experimental settings, some within-manifold perturbations required similar amounts of learning as some outside-manifold perturbations ^22^. As a result, a wide range of inputs and tasks must be considered to capture better estimates of the “true” underlying manifold, to more fully capture different neural constraints on learning.

Our context manipulations highlighted tradeoffs between interference and generalization within continual learning. Greater context signals were able to prevent overwriting by avoiding interference, particularly when learning a new within-manifold map. However, by separating the activity for different maps, context signals forced RNNs to create new activity patterns for a new within-manifold Map B instead of generalizing existing patterns that were already available for the within-manifold Map A, making learning Map B slower and more difficult (Figure 6E,F). While our implementation of context forced activity to be in very different spaces, minimizing interference but abolishing generalization, actual neural activity in different real-world contexts may be organized so as to find a compromise between interference and generalization. For example, a comparison of different but related upper limb tasks indicated that their underlying patterns were shifted with respect to each other but these tasks still shared dynamical and geometric features^49^. How the brain accomplishes such a compromise with different contexts to promote continual learning would be an interesting avenue for future work.

### Neural mechanisms for uniform shifts and memory traces

While uniform shifts and memory traces were consistently present in our RNNs without additional context inputs, we did not explicitly model any mechanisms to maintain information about different tasks in the same activity. Simply through sequential learning through gradient descent, the RNNs were pushed towards solutions that accommodated information about both tasks, rather than reverting to the original state before the new task was learned. Without explicit mechanisms, the magnitude of uniform shifts and memory traces observed may serve as a baseline minimum. While we have shown that this baseline is affected by many factors like the initial behavioral error, the brain may implement additional mechanisms that might organize these neural activity changes in a way that further fosters continual learning.

Within the machine learning literature, many neuroscience-inspired methods exist that try to encourage continual learning without catastrophic forgetting of previous tasks, including methods that regulate levels of synaptic plasticity to protect existing knowledge ^50,51^, push activity changes to be in orthogonal subspaces in the population space ^52,53^, incorporate context to separate activity ^54,55^, and use complementary learning systems that promote replay and consolidation^56^. By implementing different methods, contrasting their effects on uniform shifts and memory traces, and comparing them to experimental results, we may be able to pinpoint specific ways by which the brain may accommodate different neural activity patterns underpinning different behaviors. Since these motor memory signatures are directly experimentally measurable, they have the potential to serve as markers, enabling us to disentangle and distinguish between different types of learning.

## Conclusion

Animals build an increasingly rich behavioral repertoire by acquiring different skills during development and a lifetime of learning. Yet, how the brain accommodates novel activity patterns necessary for the generation of these new behaviors remains an open question. Recent experimental work in monkeys has proposed two potential motor cortical signatures of motor memories underpinning learning. Here, we leveraged the full access afforded by RNN models to simulate this continual learning process and systematically examine how properties of these neural signatures relate to the formation of a motor memory. We found that these signatures emerge naturally as a byproduct of sequential learning, and that their relationship with motor memories is complex: their correlation depends on the details of the perturbation that causes learning and even on whether this perturbation is explicitly cued, as expected from recent behavioral work in humans. Thus, our work identifies fundamental challenges to define a clear relationship between neural activity changes and the formation of motor memories, but also provides insights and metrics to interpret future experimental results.

## Methods

### Tasks

We trained RNNs to perform a standard delayed center-out reach task, which is commonly used in experimental settings to examine motor control. In experiments, the subject would control a cursor on a computer screen. The cursor starts in the center of a circle, and the subject has to reach one of eight possible targets evenly spaced around the circle. The subject is shown which target to move to with a target cue, but they have to delay movement until a later go cue –a design typically used to study motor planning. In our case, each trial lasted 2.0 *s*: the target cue was presented from the beginning of the trial, the go cue was presented at 0.2 *s*, and each reach lasted 1.5 *s*. We trained RNNs to produce velocities for different reaches, and the target velocities were defined by Gaussian profiles for each timestep *t* from the go cue, where each timestep equals 10 *ms* for a total of *T* = 150 time steps for each reach:

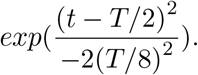

The target velocities were 0 before the go cue and after the end of the reach.

To model how experimental subjects already know how to perform different reaches prior to experiments, we first pre-trained the RNNs to produce different reaches with targets randomly distributed around a circle of 9.4 *cm* during a baseline training phase. Then, to model the experimental task, we trained RNNs to perform reaches to eight targets evenly spaced around the circle using a BCI map. The RNNs were used to model motor cortical neural activity, and the BCI map transformed unit firing rates to x and y output velocities. To examine sequential learning, we trained RNNs to use different BCI maps in blocks: first with Map A (Task A1), then with a new Map B (Task B), then back to the familiar Map A (Task A2).

### BCI maps

Map A consisted of two transformations. First, a projection matrix **C** was used to project the firing rates from all *n* = 400 units onto an 8D space, or manifold, that captured the majority of the variance in the activity. This “intuitive manifold” was determined by performing principal component analysis (PCA) on the firing rates at the end of baseline training concatenated over trials. PCA finds *n* orthogonal basis vectors (principal components or PCs) that maximally capture the variance in the population activity, sorted by their corresponding amount of variance explained. We kept the eight leading PCs, which captured more than 80% of the variance in our network activity. Second, a decoder matrix **D** was used to project activity from the manifold to a two-dimensional space indicating the output velocities. The decoder was determined offline by performing linear regression of the produced velocities on the principal components. The full map was then given by:

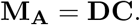

Map B was either a within-manifold perturbation or an outside-manifold perturbation of Map A. For within-manifold perturbations, the transformation from the manifold to the output velocities was disrupted with a permutation matrix *η*_**wm**_ inserted between the first and second linear transformations:

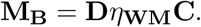

To examine the effects of perturbing different dimensions of the manifold, we examined within-manifold perturbations that permuted only the “high-variance” (first four) dimensions, only the “low-variance” (last four) dimensions, or all of the dimensions. In Figures 2-4, S1, S3, and S11, we examined the effects of 10 maps for random permutations of all dimensions. In Figure S2B-D and S5, we examined 24 maps each from all possible permutations (4! = 24) of the “high-variance” and “low-variance” dimensions, and 10 maps for random permutations of all dimensions.

For outside-manifold perturbations, the transformation from the firing rates to the manifold was disrupted with a permutation matrix *η*_**OM**_ inserted before the first linear transformation:

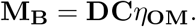

Note that *η*_*W M*_ is a 8×8 permutation matrix such that there are 8 factorial (40,320) possible *η*_*W M*_ s, while *η*_*OM*_ is a 400×400 permutation matrix such that there are 400 factorial (¿10868) possibilities. To equalize the dimensions of the search space for within- and outside-manifold perturbations, we followed methods similar to Sadtler et al. 2014. In short, we separated the 400 units into 8 groups and only considered outside-manifold perturbations where we permuted these groups, keeping all units in the same group together. We then considered all within-manifold maps and this limited set of outside-manifold maps as candidate Map Bs. Since we saw that the memory trace was highly correlated with the Task A1→Map B output MSE (Figure 4B), in Figures 5, 6, S2E-G, and S6-S10, we controlled for this factor by calculating the Task A1→Map B output MSE for all candidate Map Bs and choosing the 10 within- and 10 outside-manifold maps with MSEs closest to the median of all values.

### Neural network model

#### Network architecture

The model dynamics were given by:

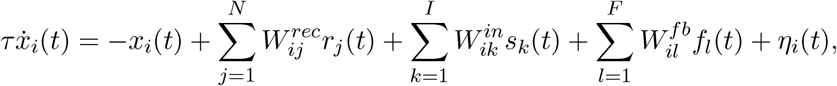

where *x*_*i*_ was the hidden state of the *i*th unit and *r*_*i*_ was the corresponding firing rate following *tanh* activation of *x*_*i*_. The network had *N* = 400 units, *I* inputs, and *F* = 2 feedback signals. The time constant *τ* was set to 0.05 *s*, the integration time step *dt* was set to 0.01 *s*, and the noise *η* was randomly sampled from the Gaussian distribution 𝒩 (0, 0.1) for each time step. The initial states **x**_*t*=0_ were sampled from the uniform distribution 𝒰 (−0.1, 0.1). The network was fully recurrently connected, with the recurrent weights **W**^**rec**^ initially sampled from the Gaussian distribution 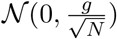, where *g* = 1.2. The time-dependent stimulus inputs **s** (specified below) were fed into the network, with input weights **W**^**in**^. The delayed feedback **f** was equivalent to the two-dimensional output velocities **v**, but delayed by 120 ms, with feedback weights **W**^**fb**^. The output *v* corresponded to *x* and *y* velocities of motor trajectories, and they were read out via the linear map:

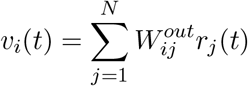

with the output weights **W**^**out**^. **W**^**in**^, **W**^**fb**^, and **W**^**out**^ all initially sampled from the uniform distribution 𝒰 (−0.1, 0.1).

To model networks with and without context, two types of sustained inputs **s** were used. For the networks without context, **s** was four-dimensional and consisted of a one-dimensional hold signal, a two-dimensional target signal (2 *cos θ*^*target*^, 2 *sinθ*^*target*^) that specified the reaching direction *θ*^*target*^ of the target, and a one-dimensional length signal that specified the normalized length of the reach. All reaches were the same length so they were kept at 2, with the same magnitude as the target and hold signals. For the networks with context, **s** was six-dimensional, with the additional dimensions consisting of a two-dimensional one-hot encoded context signal. Context A (denoted by the vector (2, 0)) was used for baseline training, Task A1, and Task A2 while Context B (denoted by (0, 2)) was used for Task B1 and Task B2. To examine the effects of different context magnitudes, we multiplied these vectors by different factors: 0.5, 1, 2, 4, 6, 8, 10, 12, 14, and 16. For both types of inputs, the hold signal started at 2 and went to 0 at the go cue while the target, length, and context signals were presented throughout the trial. While all weights (**W**^**in**^, **W**^**rec**^, **W**^**fb**^, **W**^**out**^) were initialized at the beginning of baseline training, the input weights specific for Context B were also reinitialized at the beginning of Task A1 since the weights otherwise decreased dramatically towards 0 during baseline training when Context B was not in use.

To model upstream brain regions, we added an additional RNN module similar to Feulner et al^57^. This module received sustained inputs as described above, and was fully connected to the downstream module.

#### Model training

Networks were optimized to generate velocities of synthetic center-out reaches. To model baseline learning, we pretrained the networks on reaches with targets randomly distributed around a circle of 8 cm. During baseline training, networks had to learn all the weights (**W**^**in**^, **W**^**rec**^, **W**^**fb**^, **W**^**out**^) using the Adam optimizer with an initial learning rate *l* = 10^−4^, first moment estimates decay rate *β*_1_ = 0.9, second moment estimates decay rate *β*_2_ = 0.999, and epsilon *ϵ* = 1*e* − 8. Baseline training was implemented with 500 training trials and a batch size *B* = 64. Then, to model sequential learning, we trained the networks to control different BCI maps in sequential blocks: first with Map A (Task A1), then with a new Map B (Task B), then back to the familiar Map A (Task A2). During each block, networks had to learn **W**^**in**^, **W**^**rec**^, and **W**^**fb**^, but the output weights **W**^**out**^ were fixed as either Map A (**M**_**A**_) or Map B (**M**_**B**_). The weights were learned using stochastic gradient descent with a fixed learning rate *l* = 5^−3^. We used a faster learning rate during sequential learning to model faster short-term learning compared to long-term baseline learning. Each block was implemented with 300 training trials and a batch size *B* = 64. To examine the effects of learning different tasks, we trained networks separately on different Map Bs following Task A1 (see *Tasks* for delineation of different Map Bs used).

To examine the effects of using the intuitive manifold as the familiar map, we also trained networks on the same pairs of Map As and Map Bs, but we used the Map Bs as the familiar map instead; in other words, training proceeded from baseline training, to Map B, to a new Map A, then back to the familiar Map B. To examine the effects of simultaneous rather than sequential learning, we simultaneously trained networks on the same pairs of maps during baseline learning following network initialization. All training configurations were performed on eight different networks initialized from different random seeds.

The loss *L* was the mean-squared error between the two-dimensional output and target velocities over each time step *t*, with the total number of time steps *T* = 200:

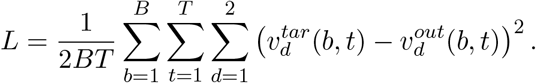

To produce network activity that aligned more closely to experimentally measured neural activity^28,29^, we added L2 regularization terms for the activity rates and network weights in the overall loss function LR used for optimization:

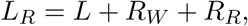

where

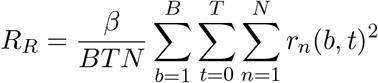

and

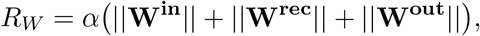

where *β* = 0.2 and *α* = 0.001. Note that the loss recorded in the main text was the loss *L* before regularization. We clipped the gradient norm at 0.2 before applying the optimization step. Following training, we evaluated the networks at each block using testing data for 320 trials (40 per target).

### Data analysis

#### Distance between population activity means

To assess how learning a new task changes activity for a familiar task, we examined how network population activity changed from Task A1 to Task A2. We obtained the population activity by projecting network activity from each of the three task blocks into the eight-dimensional manifold. To compare our results with experimental results, we followed the methods outlined for the experimental analyses in Losey et al. 2024^18^. In brief, we calculated the Mahalanobis distances between the population activity means for Task A1 and Task A2 for each target, using the covariance of Task A1 activity per target. As a control, we recalculated the Mahalanobis distances with shuffled task labels. To further demonstrate that activity is distinctly different between the tasks, we used linear discriminant analysis with five-fold cross-validation to classify the three task blocks using only the preparatory activity (first 200 *ms* of each trial).

#### Uniform shifts

To measure uniform shifts in the preparatory space, we followed the methods outlined in Sun et al. ^11^ In short, we examined population activity only in the preparatory period of each trial, which lasted 200 *ms* from trial start to the go cue. To estimate the manifold in the preparatory space, we applied PCA to the preparatory activity at the end of baseline training, then projected preparatory unit activity from all three task blocks into this manifold to get the preparatory population activity. We defined the uniform shift axis as the vector connecting the centroids of the time-averaged, trial-averaged population activity for all targets from Task A1 to Task B. We wanted to make sure that the uniform shift was not capturing changes due to behavioral output differences, so we performed targeted dimensionality reduction (TDR) on the preparatory activity and RNN outputs from Task A1 to identify a neural subspace that was predictive of behavioral output (velocities) in the upcoming reach, and we orthogonalized the uniform shift axis against the TDR axes. For each target, we projected the time-averaged, trial-averaged population activity onto the orthogonalized uniform shift axis to quantify the uniform shift per target. We obtained the normalized shifts by normalizing by the length of the target-averaged uniform shift from Task A1 to Task B. Thus, the average normalized uniform shift for all targets from Task A1 to Task B would be 1, and uniform shifts are sustained if they remain positive and close to 1 from Task A1 to Task A2.

#### Memory traces

To measure how learning a new task can leave a memory trace in the activity of a familiar task, we examined how activity patterns for a familiar task may become more useful for a new task after learning the new task. Specifically, we examined how activity patterns in Task A2 may be more appropriate for Map B than the patterns of Task A1. We projected activity from Task A1 and Task A2 into Map B offline and quantified the memory trace *m* as the increase in performance, or decrease in MSE, from Task A1 to Task A2:

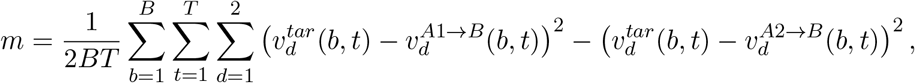

where

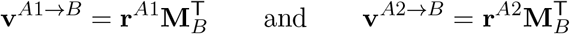

are the output velocities when Task A1 and Task A2 activity are projected into Map B, respectively.

#### Task properties

To understand how properties regarding the relationship between tasks may affect the magnitude of the memory trace, we examined two measures that quantified the relationship between Map A and Map B: the principal angles and the variance accounted for (VAF) between the maps.

Principal angles measure the relative alignment between two *m*-dimensional subspaces in an *n* dimensional space (*m < n*) by quantifying the *m* angles between pairs of basis vectors in the subspaces that minimize the angles between them. We followed the methods from Bjorck and Golub^58^ to calculate the principal angles: given the maps **M**_**A**_ and **M**_**B**_, we perform QR decomposition to obtain orthonormal bases **Q**_**A**_ and **Q**_**B**_. We then construct the inner product of these bases and perform singular value decomposition to obtain:

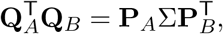

where **P**_*A*_ and **P**_*B*_ specify the directions that minimize the principal angles and **Σ** is a diagonal matrix whose entries are the ranked cosines of the principal angles.

Variance accounted for (VAF) measures how much the variance associated with a given task can be accounted for when projected into the map for a different task. We computed the variance when Task A1 activity was projected into Map A and the variance when it was projected into the new bases for Map B found using principal angles:

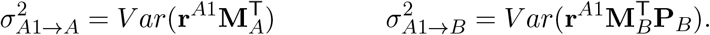

VAF was the ratio of these two variances:

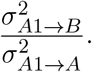

To understand how the magnitude of the memory traces relate to the initial behavioral disruption imposed by a new task, we measured the initial error using the Task A1→Map B output MSE and the Task A1→Map B angular error. The Task A1→Map B output MSE has been described above, whereas the Task A1→Map B angular error was computed based on the difference between the angle to the target and the mean angle for the entire reach during movement.

To understand how learning a new task changed the activity with respect to the manifold, we calculated the manifold overlap^59^ of Task B1 activity **r**^**B1**^ with the original intuitive manifold. We first calculated the covariance matrix **S** of **r**^**B1**^ and projected it into **C**, the first eight principal components that describe the intuitive manifold. To quantify the variance explained by the projection, we divided the trace of the projection by the trace of the corresponding covariance matrix:

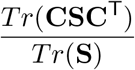

#### Control analysis

To verify that memory traces of Map B arose specifically due to the process of learning Task B, rather than due to changes that may arise from other processes, we ran two controls. First, we reset the weights learned in Task B (**W**^**in**^, **W**^**fb**^, and **W**^**rec**^) to those at the end of Task A1 before Task A2 to check if memory traces may arise due to generally more learning of Map A. Second, we randomly redistributed the weight changes incurred during Task B before Task A2 to check if memory traces may arise due to random weight changes of the same magnitude that are unrelated to learning Map B.

## Code availability

All analyses were implemented using custom python code (Python 3.8) and open-source software. All the figures are reproducible by running Jupyter notebooks. Code to reproduce all the results is openly available on GitHub at https://github.com/JoannaChang/motor_memory.

## Acknowledgements

J.C.C. received funding from the Wellcome Trust (grant 108908/Z/15/Z). C.C received funding from the BBSRC (BB/N013956/1 and BB/N019008/1), the EPSRC (EP/R035806/1), the Wellcome Trust (200790/Z/16/Z), and Simons Foundation (564408). J.A.G. received funding from the EPSRC (EP/T020970/1) and the European Research Council (ERC-2020-StG-949660). The funders had no role in study design, data collection and analysis, decision to publish, or preparation of the manuscript.

## Author Contributions

J.C.C., J.A.G. and C.C. devised the project. J.C.C. ran simulations, analysed data and generated figures. J.C.C., C.C. and J.A.G. interpreted the data. J.C.C., C.C. and J.A.G. wrote the manuscript. All authors discussed and edited the manuscript. J.A.G. and C.C. jointly supervised the work.

## Supplementary Figures

**Supplementary Figure S1:**
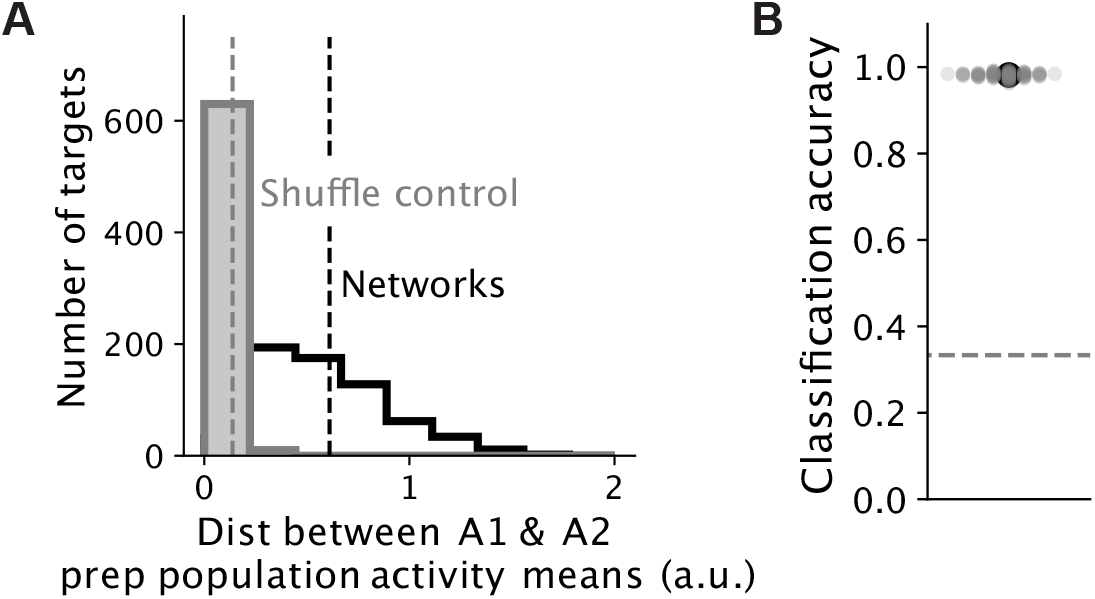
Neural activity changes from Task A1 to Task A2. **A)** Mahalanobis distance between population activity means for Task A1 and Task A2 for each target during preparation (first 200 ms of each trial, black), and for controls where the task labels were shuffled (gray). Dotted lines, means across 10 different maps for each random seed (*n*=8 random seeds). **B)** Classification accuracy for classifying tasks (Task A1, Task B1, Task B2) from neural activity during preparation using linear discriminant analysis. Large circle, mean across maps and random seeds; small circles, accuracy for each map for each random seed; colors, different random seeds; dotted line, chance accuracy level.

**Supplementary Figure S2:**
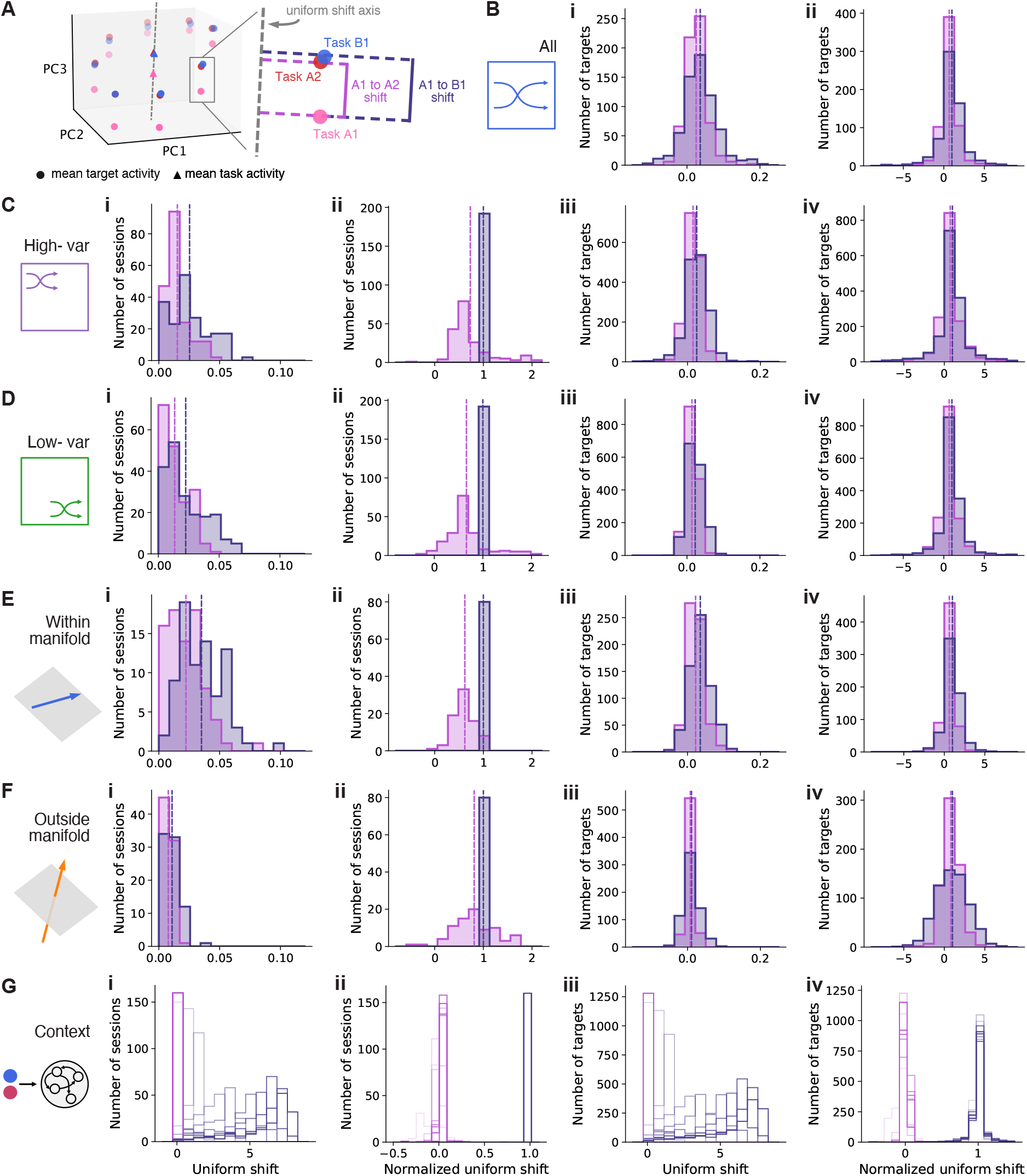
Uniform shifts for different simulations. **A)** Uniform shifts can be quantified per target for each session rather than averaged across all targets for each session as in Fig. 3B. To quantify uniform shifts per target, mean target activity (circles) was projected on the uniform shift axis determined based on mean task activity (triangles) for Task A1 and Task B1. The uniform shifts were then the resulting distances between these projected mean states. **B)** Uniform shifts for 10 different maps based on within-manifold permutations that shuffled all manifold dimensions (used in Figs 2-4). **B.i)** Uniform shifts from Task A1 to Task B1 and Task A1 to Task A2 for each target. Dotted lines, means across 10 different maps for each random seed. **B.ii)** Uniform shift per target, normalized by the uniform shift averaged across targets from Task A1 to Task B1 for each session. **C)** Uniform shifts for 24 different maps based on within-manifold perturbations that shuffled the first four high-variance dimensions (used in Fig 4). **C.i)** Uniform shift averaged across targets (as in Fig. 3B). **C.ii)** Normalized uniform shift averaged across targets (as in Fig. 3C). **C.iii-iv)** Same as B.i-ii but for high-variance maps. **D)** Same as C but for 24 different maps based on within-manifold perturbations that shuffled the last four low-variance dimensions (used in Fig S6). **E)** Same as C but for 10 different within-manifold perturbations with the same initial output MSE (used in Fig 5). **F)** Same as C but for 10 different outside-manifold perturbations with the same initial output MSE (used in Fig 5). **G)** Same as C but for 10 outside-manifold and 10 within-manifold perturbations with the same initial output MSE for networks with different context magnitudes (used in Fig 6). Opacities, different context magnitudes (most transparent for smallest magnitude).

**Supplementary Figure S3:**
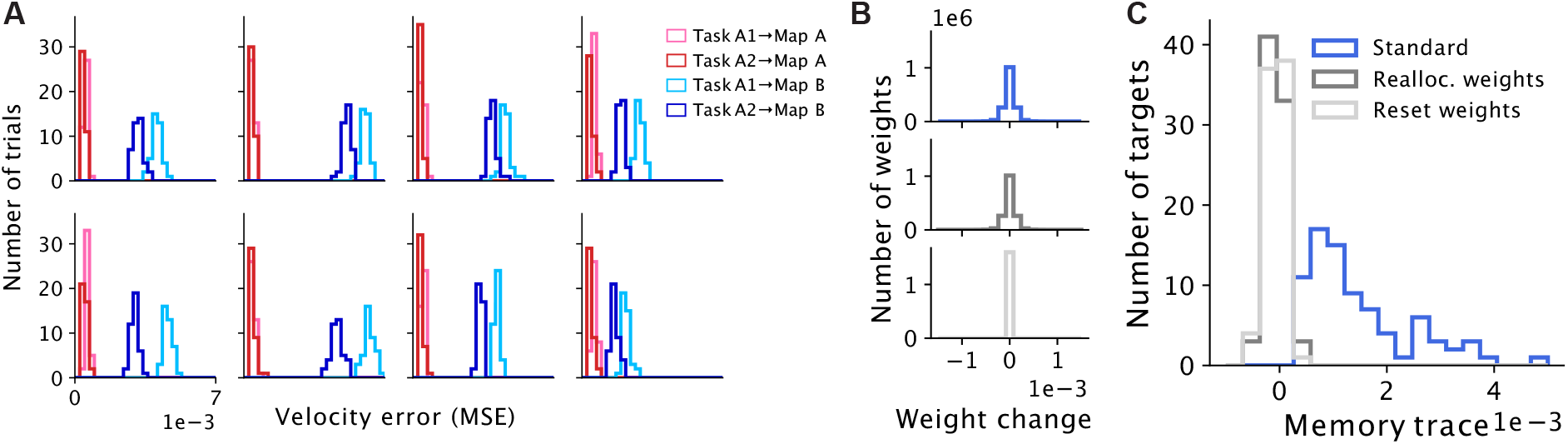
Memory traces consistently arise and are due to learning Map B. **A)** MSE between the produced and target output for Task A1 and Task A2 activity when projected into Map A and Map B for each target in an example session (same session as Fig 3E). **B-C)** To show that memory traces arise specifically due to changes incurred from learning Map B, networks were trained on Task A2 without altered weights (standard, blue), with weight changes from Task B randomly reallocated before Task A2 (dark gray), or with weights reset before Task A2 to those at the end of Task A1 (light gray). **B)** Weight changes from end of Task A1 to beginning of Task A2. **C)** Memory traces for each target for each session for 10 different maps for each random seed (*n*=8 random seeds) after the weight manipulations outlined in B.

**Supplementary Figure S4:**
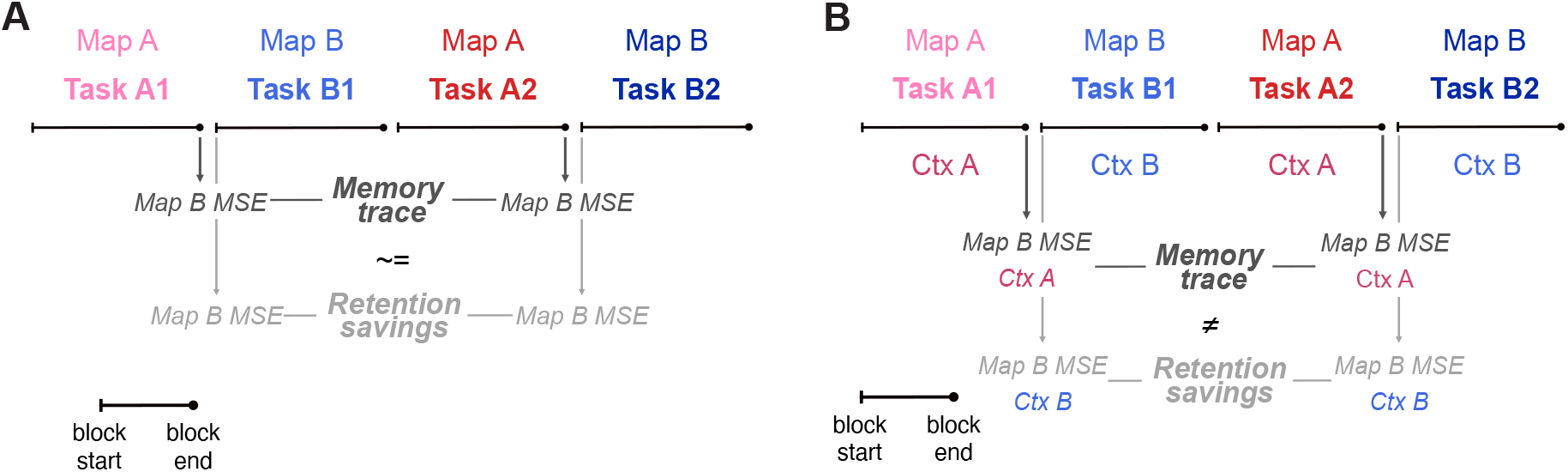
Relationship between memory trace and retention savings. A) Schematic for how memory trace and retention savings are measured based on MSEs at different stages of learning without context cues. Note that the memory trace is closely related to retention savings. The memory trace is defined as the increase in performance from Task A1→Map B to Task A2→Map B while retention savings is defined as the increase in performance from the beginning of Task B1→Map B to the beginning of Task B2→Map B. Without context cues, the network dynamics are identical at the end of Task A1 (Task A2) compared to the beginning of Task B1 (Task B2), other than minimal contributions from feedback, such that the memory trace is equivalent to retention savings, other than the contributions from feedback. **B)** Schematic for how memory trace and retention savings are measured based on MSEs at different stages of learning with context cues. Note that memory trace is not equivalent to retention savings when context cues are added because the network dynamics at the end of Task A1 (Task A2) are not identical to those at the beginning of Task B1 (Task B2).

**Supplementary Figure S5:**
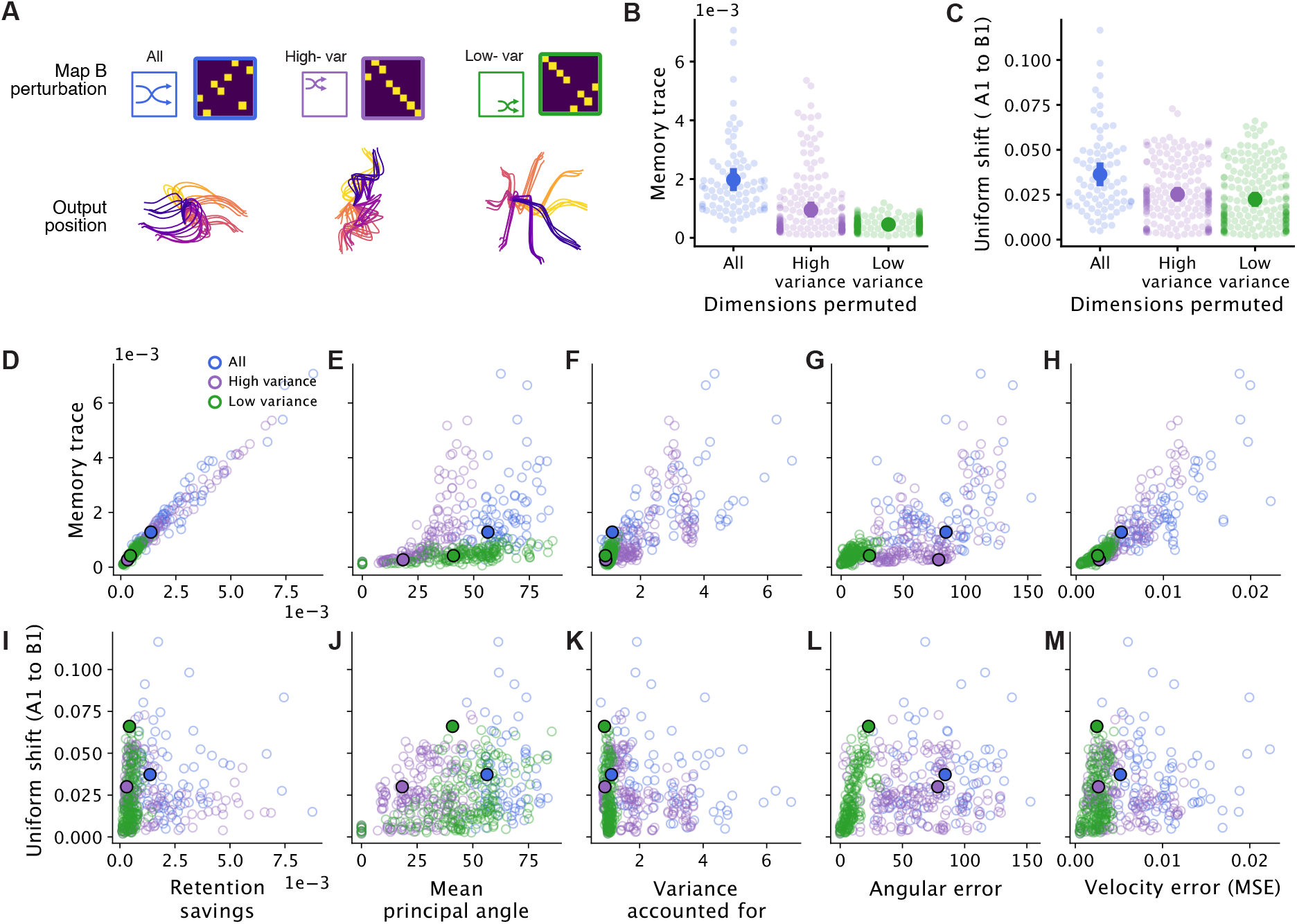
The magnitude of memory traces and uniform shifts compared to different measures. **A**) Networks were trained on different Map Bs representing within-manifold perturbations made from permutations in “all” manifold dimensions (blue, *n*=10 maps), only the first four “high-variance” dimensions (purple, *n*=24 maps), or only the last four “low-variance” dimensions (green, *n*=24 maps). Top: Permutation matrices for example maps for each category. Bottom: Output positions when Task A1 activity is projected into the new maps before learning in Task B1. **B)** Mean memory trace across targets. Big circle and error bars, means and 95% confidence intervals with bootstrapping across different maps for each random seed (n = 8 random seeds); small circles, different maps for different random seeds. **C)** Uniform shift from Task A1 to Task B1 across targets. **D-H)** Mean memory trace across targets compared to different measures: from left to right, retention savings, mean principal angle between Map A and Map B, variance accounted for (VAF) in Task A1 activity by Map B, Task A1**→**Map B angular error, and Task A1**→**Map B velocity MSE (Methods). Open circles, different maps for different random seeds; closed circles, examples for each permutation type shared across D-M. **I-M)** Uniform shift from Task A1 to Task B1 across targets compared to different measures.

**Supplementary Figure S6:**
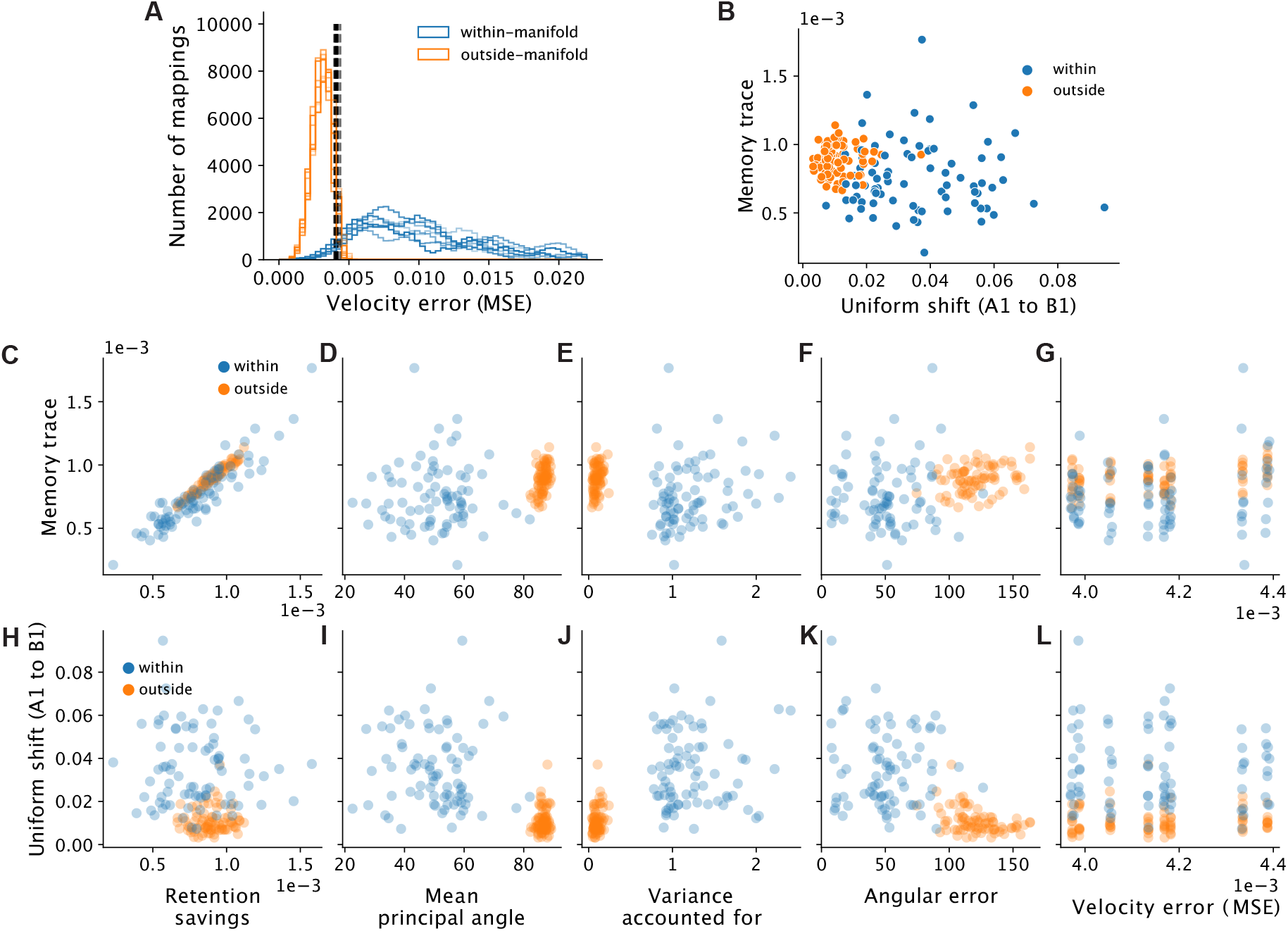
Relationship between memory trace and uniform shift and different factors once initial MSE is fixed. **A)** Task A1 activity was projected into thousands of candidate BCI maps representing within- and outside-manifold perturbations to obtain the initial Task A1**→**Map B MSE (Methods). Dashed lines, median for maps for each random seed. Opacities, different random seeds (*n*=8 random seeds). The 10 maps for each type of perturbation with MSEs closest to the median MSE across all maps for each seed were chosen for training. B) Uniform shift from Task A1 to Task B1 across targets compared to the mean memory trace across targets. Circles, mean for each map for each random seed. C-G) Mean memory trace across targets compared to the retention savings. **C-G)** Mean memory trace across targets compared to different measures: from left to right, retention savings, mean principal angle between Map A and Map B, variance accounted for (VAF) in Task A1 activity by Map B, Task A1**→**Map B angular error, and Task A1**→**Map B velocity MSE (Methods). Circles, different maps for different random seeds. **H-L)** Uniform shift from Task A1 to Task B1 across targets compared to different measures.

**Supplementary Figure S7:**
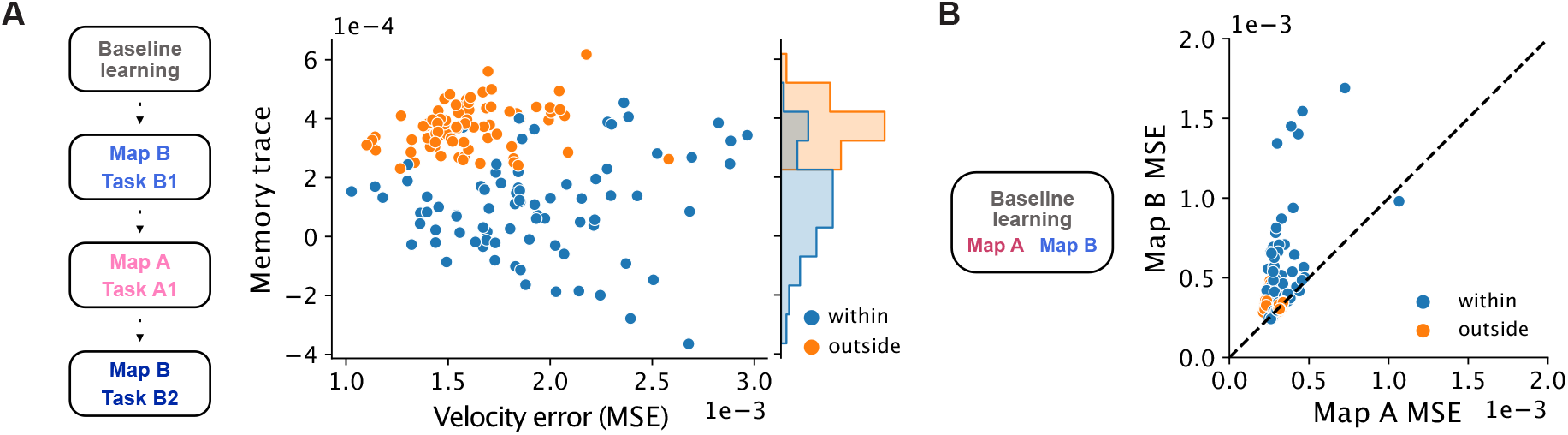
Learning within- and outside-manifold maps under different training regimes. **A)** Networks were trained with the same Map A and Map B pairings as in the main text (Fig 5), but the within- or outside perturbation (Map B) was used as the familiar map while the intuitive map (Map A) was used as the new map, such that learning proceeded from baseline learning, to Map B, to Map A, and back to Map B. Memory trace compared to Task A1**→**Map B MSE. Circles, mean for each map (*n*=10 maps per perturbation type) for each random seed (*n*=8 random seeds). **B)** Networks were trained with the same Map A and Map B pairings, but they were trained simultaneously during baseline learning after network initialization. Map A MSE compared to Map B MSE after baseline learning. Circles, mean for each map for each random seed; dashed line, identity line. Note that Map B had worse performance than Map A, especially for within-manifold perturbations.

**Supplementary Figure S8:**
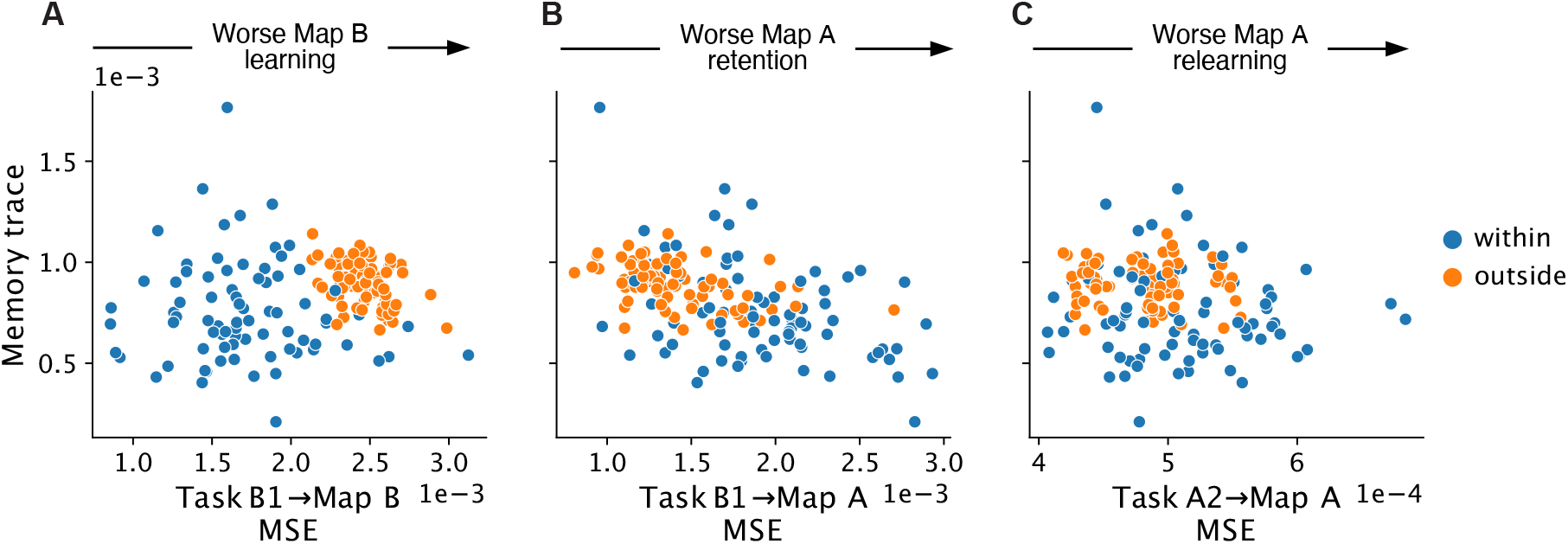
Larger memory traces for outside-manifold learning are not due to performance tradeoffs. **A)** Memory traces compared to Task B1**→**Map B MSE. Greater memory traces for outside-manifold perturbations were not due to better initial learning of Map B in Task B1. Circles, mean for each map (*n*=10 maps per perturbation type) for each random seed (*n*=8 random seeds). **B)** Memory traces compared to Task B1**→**Map A MSE. Greater memory traces for outside-manifold perturbations were not due to worse retention of Map A in Task B1. **C)** Memory traces compared to Task A2**→**Map A MSE. Greater memory traces for outside-manifold perturbations were not due to worse relearning of Map A in Task A2.

**Supplementary Figure S9:**
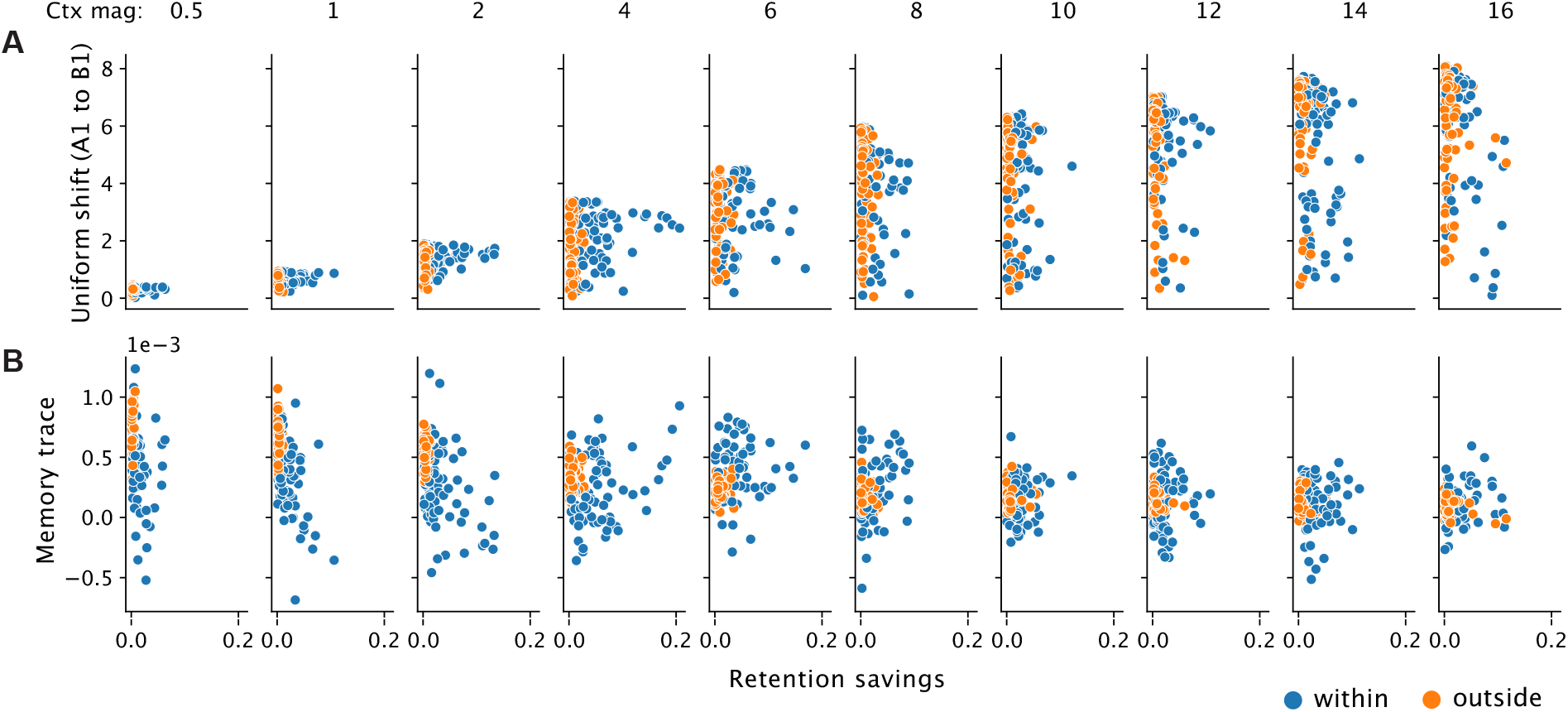
Context disrupts the correlation between memory traces and savings. **A)** Uniform shift from Task A1 to Task B1 across targets compared to mean retention savings for different context magnitudes. Circles, mean for each map (*n*=10 maps per perturbation type) for each random seed (*n*=8 random seeds). **B)** Mean memory trace across targets compared to mean retention savings for different context magnitudes.

**Supplementary Figure S10:**
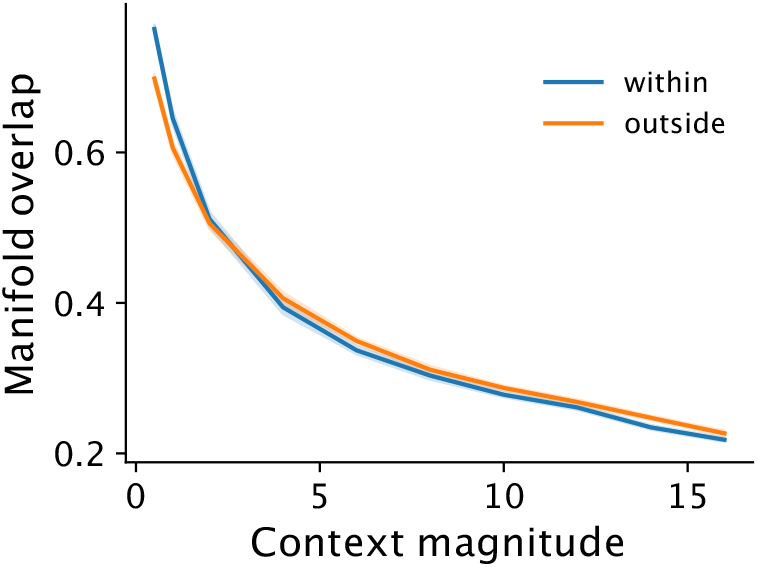
Manifold overlap for different context magnitudes. Manifold overlap between the activity after learning the new map in Task B1 and the original intuitive manifold for within- and outside-manifold perturbations. Line and shaded surfaces, mean and 95% confidence interval across 10 maps for each perturbation across networks of different seeds (*n*=8 random seeds).

**Supplementary Figure S11:**
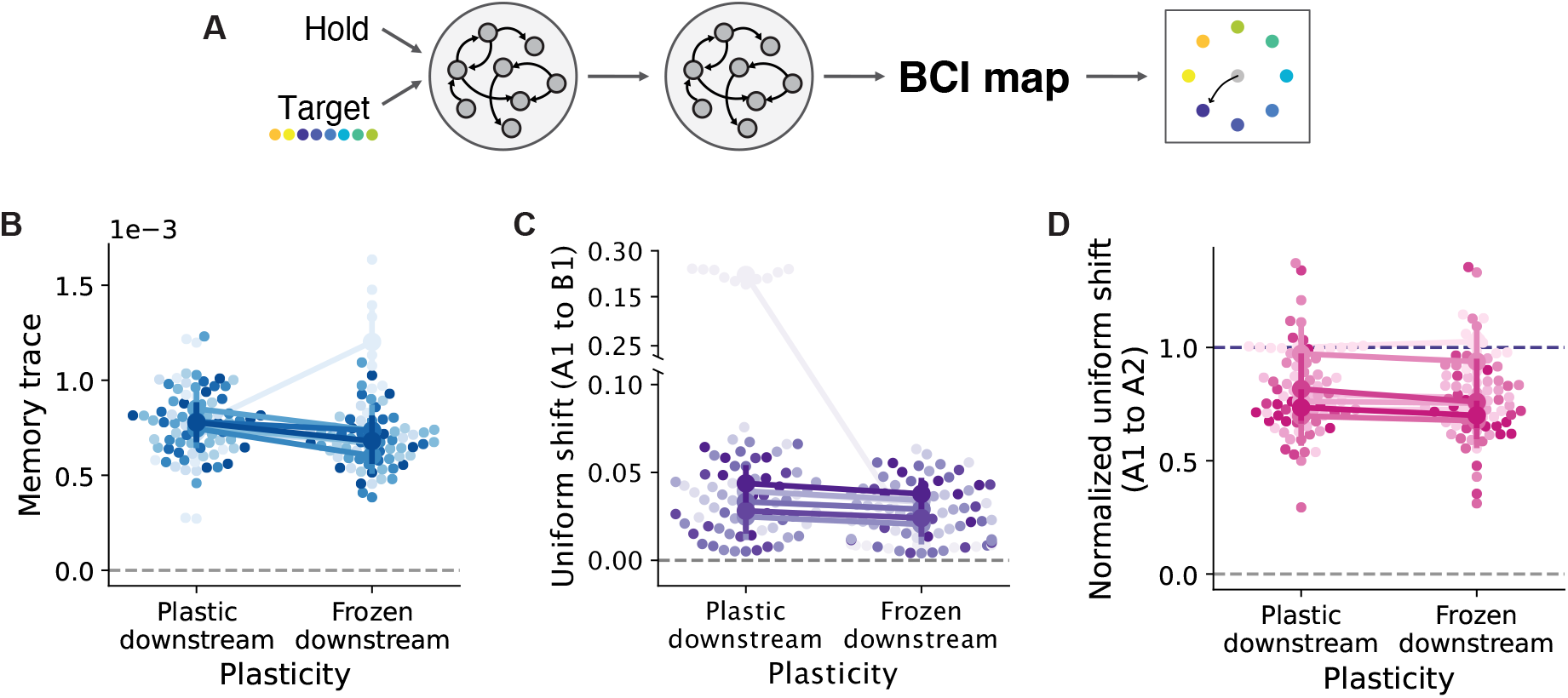
Memory traces and sustained uniform shifts arise when learning is confined to upstream of motor cortex. **A)** Schematic for network models with an additional RNN module added upstream. Activity from the downstream module is directly used to control the BCI, similar to how activity from the motor cortex is often used to control BCIs. The additional upstream module models brain regions upstream of the motor cortex. We examined how memory traces and uniform shifts differed when learning was restricted to the upstream module (frozen downstream weights) compared to when learning was allowed in the downstream module (plastic downstream weights). **B)** Mean memory trace across targets. Big circle and error bars, means and 95% confidence intervals with bootstrapping for 10 different maps for each random seed (*n*=8 random seeds). Small circles, different maps; colors, different random seeds. Note that memory traces were consistently greater than zero (grey dotted line), even when learning is restricted upstream. **C)** Uniform shift from Task A1 to Task B1 across targets. **D)** Uniform shift from Task A1 to Task A2 across targets, normalized to the uniform shift from Task A1 to Task B1. Note that the uniform shifts were sustained since they were consistently greater than zero (grey dotted line) and close to the shift from Task A1 to Task B (purple dotted line).

## Footnotes

1 Recall that the memory trace is quantified as the increase in performance, or decrease in MSE, from Task A1**→**Map B to Task A2**→**Map B, so it is inversely proportional to the Task A2**→**Map B MSE when we have controlled for the initial output error, the Task A1**→**Map B MSE. We can thus use the trends in Task A2**→**Map B MSE to examine the trends in memory traces in Figure 6E and F).

